# Chemoproteomics reveals immunogenic and tumor-associated cell surface substrates of ectokinase CK2α

**DOI:** 10.1101/2024.03.20.585970

**Authors:** Corleone S. Delaveris, Sophie Kong, Jeff Glasgow, Rita P. Loudermilk, Lisa L. Kirkemo, Fangzhu Zhao, Fernando Salangsang, Paul Phojanakong, Juan Antonio Camara Serrano, Veronica Steri, James A. Wells

## Abstract

New epitopes for immune recognition provide the basis of anticancer immunity. Due to the high concentration of extracellular adenosine triphosphate in the tumor microenvironment, we hypothesized that extracellular kinases (ectokinases) could have dysregulated activity and introduce aberrant phosphorylation sites on cell surface proteins. We engineered a cell-tethered version of the extracellular kinase CK2α, demonstrated it was active on cells under tumor-relevant conditions, and profiled its substrate scope using a chemoproteomic workflow. We then demonstrated that mice developed polyreactive antisera in response to syngeneic tumor cells that had been subjected to surface hyperphosphorylation with CK2α. Interestingly, these mice developed B cell and CD4+ T cell responses in response to these antigens but failed to develop a CD8+ T cell response. This work provides a workflow for probing the extracellular phosphoproteome and demonstrates that extracellular phosphoproteins are immunogenic even in a syngeneic system.

## Introduction

The aberrant metabolism of tumors can give rise to aberrant post-translational modifications of proteins on the tumor cell.^1–5^ Either spontaneously or with the help of immunotherapies, these modifications can be so foreign that they are recognized by the immune system leading to tumor clearance. New or over-abundant sites of protein phosphorylation (phosphoneoantigens) have been shown to be the targets of anti-tumoral immunity clinically, resulting from intracellular hyperphosphorylation leading to MHC-I display and are also targets of humoral immunity in patients.^1^ We hypothesized that the immunogenicity of aberrant phosphorylation motifs may expand to extracellular phosphoproteins.

Tumors create a distinct metabolic environment in which unique chemistries can occur due to an aberrant composition of metabolites. Notably, extracellular concentration of adenosine triphosphate (ATP) can be 10,000 to 100,000-times higher in the tumor microenvironment (up to 1 mM) compared to the interstitial space of healthy tissues.^6–8^ Extracellular ATP is a multifaceted molecule, with roles in growth signaling,^9–11^ immune modulation,^12–14^ and catalysis.^15–18^ One of the most direct roles for ATP in catalysis is as a substrate for kinases – proteins that form chemical adducts of phosphates onto other proteins. The measured intra-tumoral concentrations of extracellular ATP are just in the range of their Km values necessary for maximal enzymatic activity of many kinases (typically ranging from 1 to -100 μM).^19^

Ectokinases are a class of protein kinases with described extracellular activity that have gained increasing recognition in recent years. ^20–23^ We hypothesized that, in the context of chronically high extracellular ATP of the tumor microenvironment, these kinases may introduce immunogenic phosphoproteins on the cell surface (**Fig. 1**). We chose to investigate the ectokinase CK2α due to the established cancer-association of CK2^24–27^ and the amenability of the kinase to protein engineering for developing a proof-of-concept ectophosphoproteomic workflow.^19,28^

**Figure 1.**
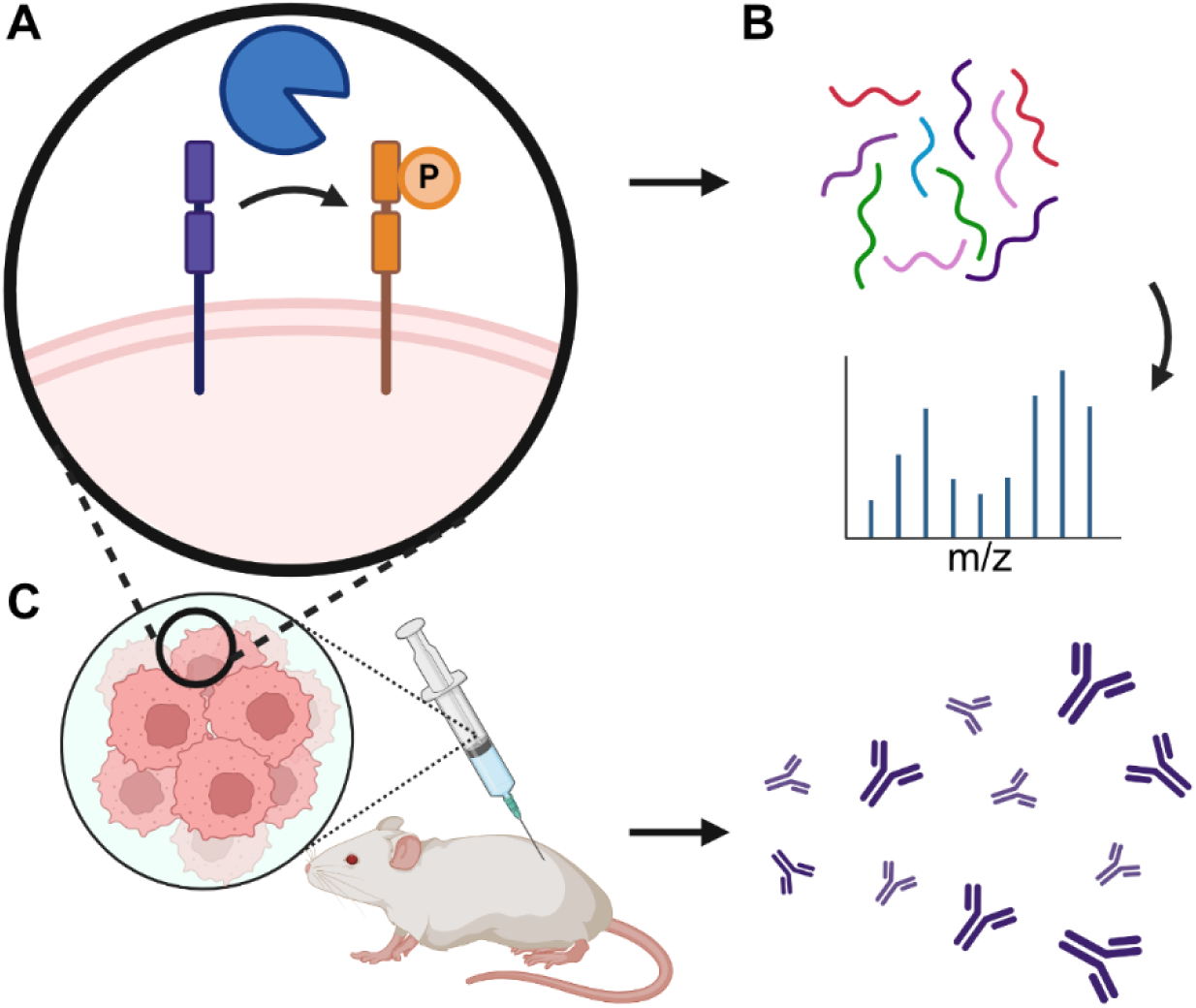
Ectokinase CK2 has cell surface substrates that elicit an immune response. **(A)** Targeting kinases to cell surfaces generates phosphorylated cell surface proteins. **(B)** Cell surface phosphoproteins are chemically analyzed by mass spectrometry. **(C)** Mice immunized with whole-cell lysates of hyperphosphorylated tumor cells generate an immune response directed against cell-surface phosphorylated proteins.

We engineered cell-tethered forms of CK2α, which were more efficient in surface phosphorylation *in vitro* than the free enzyme. We used ATP-gamma-thiophosphate (ATPγS) as a chemical reporter to probe the surface substrates of CK2α, identifying and validating substrates, including CD44, by mass spectrometry and by Western blot. We further demonstrate that this workflow can be used to interrogate endogenous CK2α using a CK2α-specific inhibitor, CX-4945. We found that the mice generated antibody responses to phosphoprotein substrates of CK2α on the surfaces of syngeneic tumor cells, hypothesizing that the phosphorylated forms of these proteins are indeed neoantigens. We characterized the immune response, identifying VH sequences specific to CK2-phosphorylated CD44, and observing that B cells and CD4+, but not CD8+, T cells responded to these neoantigens. These results highlight the importance of the extracellular phosphoproteome in anti-cancer immunity, and provide a workflow for interrogating the substrates of specific ectokinases.

## Results

### A surface tethered CK2 derivative is active on the cell surface

We chose to pursue CK2 as our lead kinase due to known activity in the extracellular space,^18,29^ ease of expression, and the established over-expression of CK2 at the protein level in breast cancer.^26,27^ To tether CK2 to the cell surface, we fused the kinase domain of CK2, CK2α, to either a HER2-binding affibody, ZHER2, (**Fig. 2A, Fig. S1**) or an Fc-knob domain for expression as a heterobifunctional knob-in-hole Fc fusion with the antigen binding domain of trastuzumab (**Fig. S2**). These targeted forms bound to the surface of hHER2-expressing EMT-6 murine breast cancer cells by flow cytometry and microscopy (**Fig. 2B, Fig. S3-4**). In these experiments, we observed some binding of CK2α to the cell surface, which has been previously proposed to be due to a tetra-lysine motif binding to heparin.^30,31^ We generated several mutants of this motif and showed that loss of positive charge abrogated cell binding (**Fig. S3**). Notably K75E and K74/75E, bound markedly less yet retained enzymatic activity. Satisfyingly, we found that the HER2 targeted construct restored binding to cells and to a much higher degree than a soluble form CK2α.

**Figure 2.**
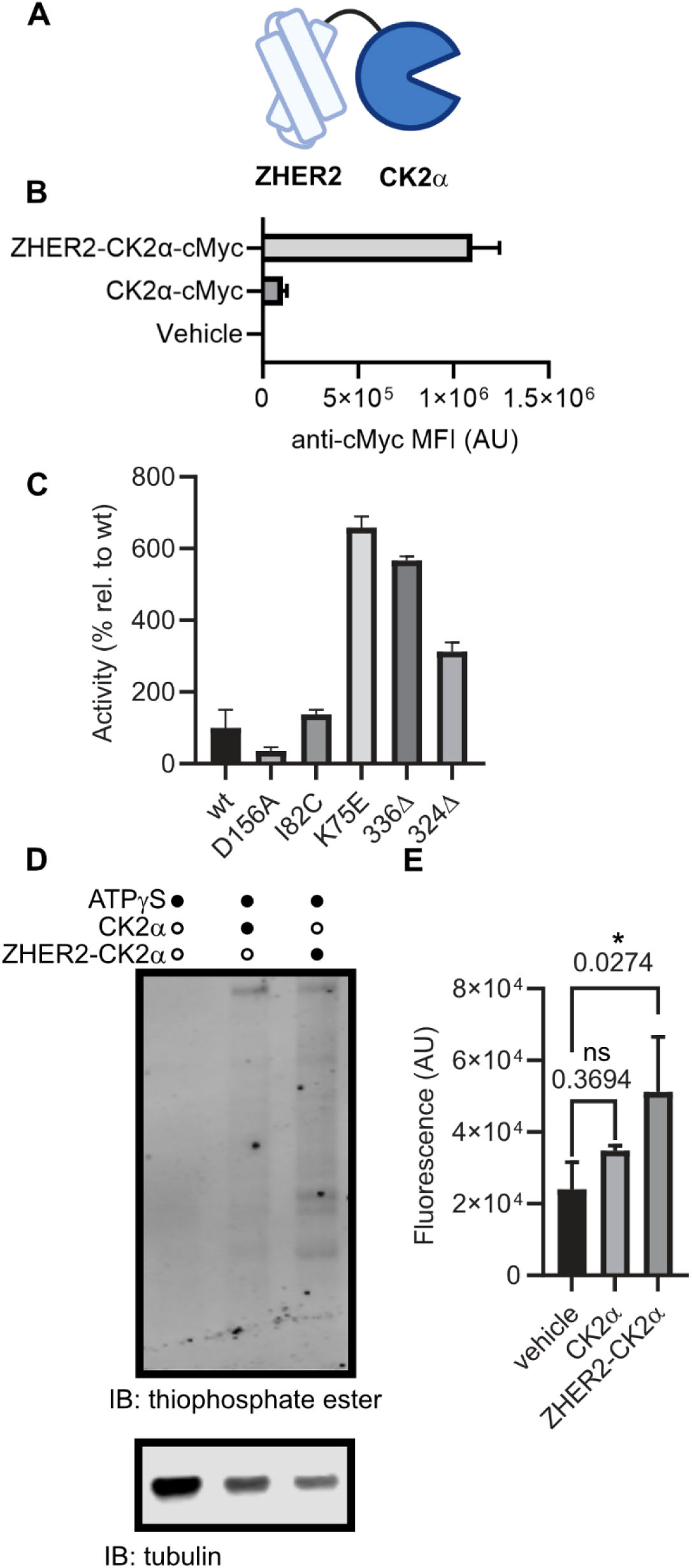
Cell-tethered CK2α is active on the cell surface under tumor-relevant conditions. **(A)** Schematic representation of kinase-fusion proteins generated. **(B)** Purified recombinant ZHER2-CK2α or CK2α proteins, C-terminally fused to a cMyc peptide tag, were incubated with hHER2-expressing EMT-6 murine breast cancer cells, and kinase binding was visualized with a fluorophore-conjugated anti-cMyc antibody by flow cytometry. **(C)** Generation of CK2α isoforms that are more stable and active. CK2α mutants were generated and assayed for activity on a model peptide substrate (RRRDDDSDDD) in tumor-relevant concentrations of ATP (100 micromolar). **(D,E)** hHER2^+^ EMT-6 cells were incubated with kinase fusion and ATP-gamma-thiophosphate in kinase reaction buffer, before being lysed, alkylated with p-nitrobenzylmesylate, separated by SDS-PAGE, and transferred to a blotting membrane. Ectophosphorylation was visualized with a thioester-specific antibody and quantitated by near-infrared fluorescence imaging. (**D**) Representative Western Blot. (**E**) Fluorescence quantitation from three independent blots. Statistics were determined by one-way ANOVA corrected for multiple comparisons. * = corrected p value < 0.05. n.s. = not significant. Corrected p-values are displayed above relevant comparisons.

To further optimize our enzyme for rapid *in vitro* labeling, we wanted to ensure that the kinase would be potently active at the concentrations of ATP relevant to the tumor microenvironment. Importantly, the Km of CK2α for ATP (∼300 µM) is in the range of what has been described for the tumor microenvironment suggesting that a kinetically tuned mutant would label faster, but not necessarily more than a kinase *in vivo*.

Therefore, we screened several previously described mutants of CK2α under tumor-relevant ATP conditions (**Fig. 2C**). Mutation of K75E both increased the activity at 100 µM ATP, as has been previously described,^19^ and decreased non-specific binding to cells. Therefore, we proceeded with the K75E mutant of CK2α moving forward. Importantly, literature has shown that the mutation K75E does not affect the substrate preference of CK2.^32,33^

We hypothesized that our tethered CK2 would be more efficient at phosphorylating the cell than an untethered version, as both we^34,35^ and others^36,37^ have observed that tethering enzymes to the cell surface increases enzymatic activity on cell surface substrates. We treated hHER2+ EMT-6 cells with a cell-tethered or soluble CK2α in the presence of a chemical reporter substrate, ATPγS, as previously utilized by Shokat and coworkers.^38^ We then lysed cells, alkylated lysates with p-nitrobenzylmesylate, and visualized kinase activity by Western blot using a highly-specific anti-thiophosphate ester antibody (**Fig. 2D-E, Fig. S5**). As we hypothesized, we observed a statistically significant increase in thiophosphate ester binding to tethered-CK2α treated cells compared to no enzyme (p = 0.046), but not for soluble CK2α (p = 0.15).

### Chemoproteomics enables selective enrichment of extracellular phosphoproteins

To define the cell surface substrate scope of CK2α, we developed an ectophosphoproteomic enrichment method (**Fig. 3A**).^38^ In this protocol, we treated hHER2+ EMT-6 cells with active or mutationally inactive cell-tethered CK2α in the presence of the cell-impermeable chemical reporter substrate ATPγS, allowing isolation of a phosphatase-resistant thiophosphorylated protein. We then lysed cells and immobilized thiophosphoproteins on iodoacetyl agarose, before eluting bound proteins with trypsin. Tryptic peptides were then identified by LC-MS (**Table S1**). We also attempted to selectively elute thiophosphoproteins by mild oxidation but were unable to confidently identify phosphopeptides.

**Figure 3.**
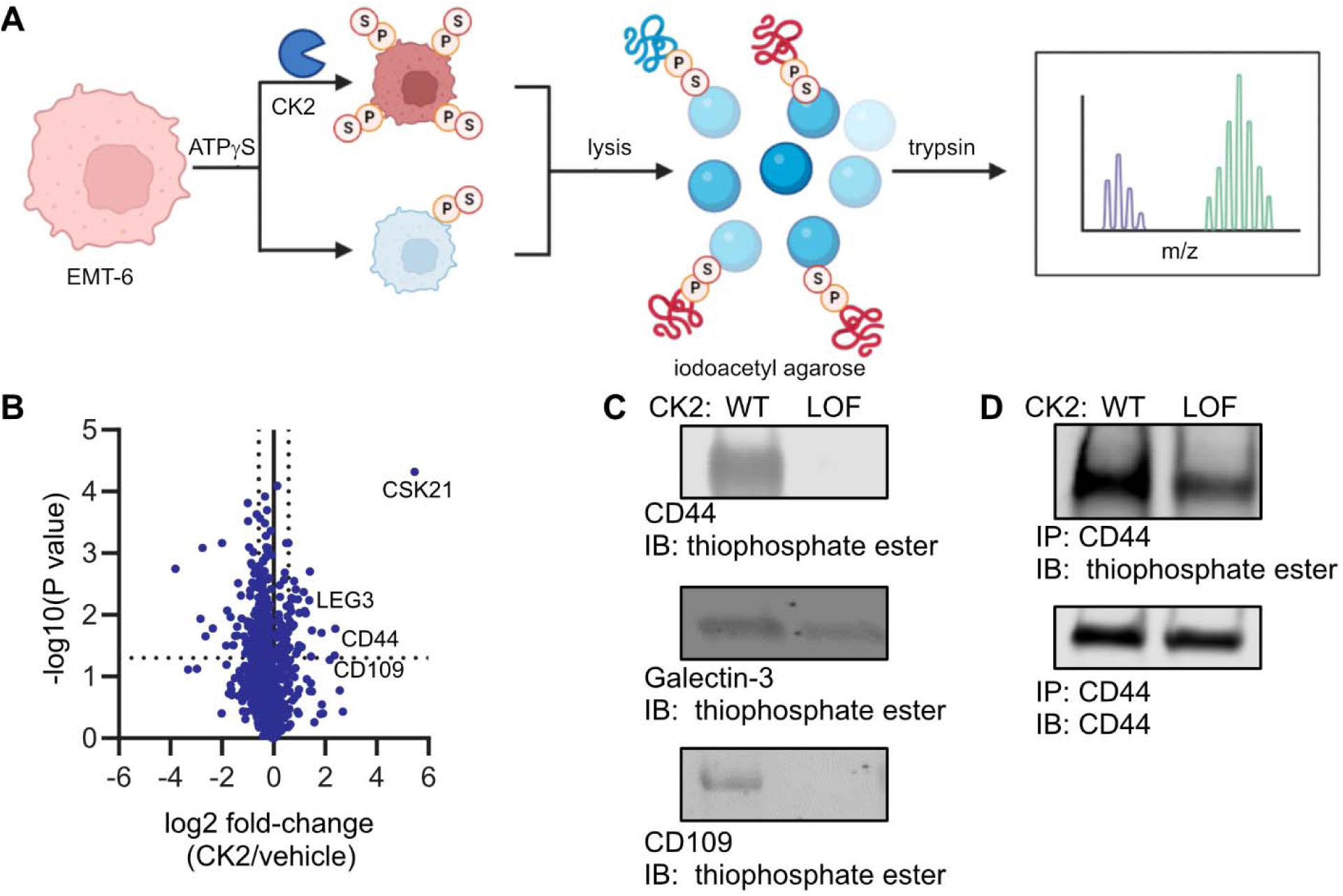
Identification of CK2 substrates using an ATP-gamma-thiophosphate (ATPγS) chemical reporter substrate using label-free quantitation based proteomics. **(A,B)** Wild-type or loss-of-function CK2α is tethered to the cell surface in the presence of ATPγS, lysed, immobilized on iodoacetyl agarose, and then tryptic eluates are analyzed by LC-MS **(B). (C)** Candidate recombinant CK2 substrates were treated *in vitro* with recombinant CK2, alkylated with p-nitrobenzylmesylate, and analyzed by Western blot with a thiophosphate ester-specific antibody. **(D)** Wild-type or loss-of-function CK2α was tethered to the surface of hHER2+ EMT-6 cells in the presence of ATPγS. Cells were lysed and CD44 was immunoprecipitated from lysates. Immunopreciptates were alkylated with p-nitrobenzylmesylate and analyzed by Western blot with a thiophosphate ester-specific antibody showing active kinase-dependent increase in CD44 phosphorylation.

We thus proceeded to use label-free quantitation (LFQ) to determine peptide abundance in replicates of cell-tethered wild-type (WT)CK2α versus a similarly-tethered loss-of-kinase-function (LOF) mutant (corresponding to D156A) (**Fig. 3B, Table S2**). We found significant enrichment of 14 proteins (>1.5 fold change, ≥ 2 peptides identified in each sample), including the tumor-associated proteins CD44,^39,40^ Galectin-3,^41^ and CD109.^42^ Not surprisingly the most significantly enriched peptides were identified from CK2α, which is known to autophosphorylate.^43^ We validated CD44, Galectin-3, and CD109 as substrates of CK2 using recombinant protein and kinase *in vitro*, visualizing the reaction by Western blot (**Fig. 3C, Fig. S6**). We further validated CD44 as a substrate on cells by treating hHER2+ EMT-6 cells with ATPγS in the presence of WT or LOF cell-tethered ZHER2-CK2α. We then probed alkylated lysates as before by Western blot and probed anti-CD44 immunoprecipitants (**Fig. 3D, Fig. S7**). We observed a marked increase in fluorescence of the anti-thiophosphate ester antibody in CD44 immunoprecipitants for the active enzyme condition compared to the loss-of-function, and comparable amounts of total immunoprecipitated CD44. Finally, we site-localized CK2α-mediated phosphorylation on the N-terminal domain of CD44 by treating recombinant human CD44 *in vitro* with soluble CK2 holoenzyme and analyzing tryptic digests by LC-MS; we identified two high confidence phosphorylation sites at S183 and S185 (**Table S3**). We additionally identified one high confidence CK2-phosphorylation site on human CD44 at T174 which sits in a canonical CK2-phosphorylation motif (**T**DDD) (**Table S4**). Collectively, these data provide strong evidence that CK2 is active on the cell surface and has CD44 among its surface substrates.

We also sought to investigate if our workflow could be expanded to probe endogenous ectokinases. To do so, we probed endogenous CK2α using a clinical CK2 inhibitor, CX-4945 (silmitasertib),^44^ in the context of the human breast cancer cell line SK-BR-3. We incubated SK-BR-3 cells in the presence of ATPγS and with either CX-4945 or an equal volume of DMSO, and the proceeded to enrich and identify phosphoproteins as before (**Table S5**). In doing so, we observed down-regulation of three proteins by LFQ (**Table S6**), all of which have reported phosphosites in a CK2α substrate motif. Furthermore, CD44 peptides were identified in the DMSO samples but not CX-4945 treated samples, preventing quantitation by LFQ. There were fewer proteins confidently identified (3 vs. 14) in the CK2α-inhibited samples compared to no treatment. Our data show this workflow can identify endogenous ectokinase substrates and that we can significantly enhance the degree and extent of substrate identification using the tethered ectokinase.

### Cell surface CK2 substrates elicit antibody and CD4+ T cell responses *in vivo*

To probe the immunogenicity of the cell surface CK2 substrates, we immunized Balb/c mice with heat-killed syngeneic hHER2+ EMT-6 tumor cells analogous to previous protocols for whole cell immunization.^45^ hHER2+ EMT-6 cells were pretreated either with our tethered-CK2α (Group 1) or vehicle (Group 2) (**Fig. 4A**). We then harvested the sera from immunized mice and measured binding to live hHER2+ EMT-6 cells treated *in vitro* with cell-tethered CK2α or vehicle by flow cytometry (**Fig. 4B, Fig. S8**). While the sera bound to all cells more than an isotype, implying some level of immunoreactivity, we observed a dramatic increase in binding to CK2-treated hHER2+ EMT-6 cells over control hHER2+ EMT-6 cells with sera from Group 1 but not Group 2. We confirmed that these were specifically CK2-substrates by engineering a bump-hole^46,47^ pair of CK2α F113G with N6-phenylethyl-ATP and validating that Group 1 sera bound specifically to cells treated with the bump-hole pair (**Fig. 4C, Fig. S9-10**).

**Figure 4.**
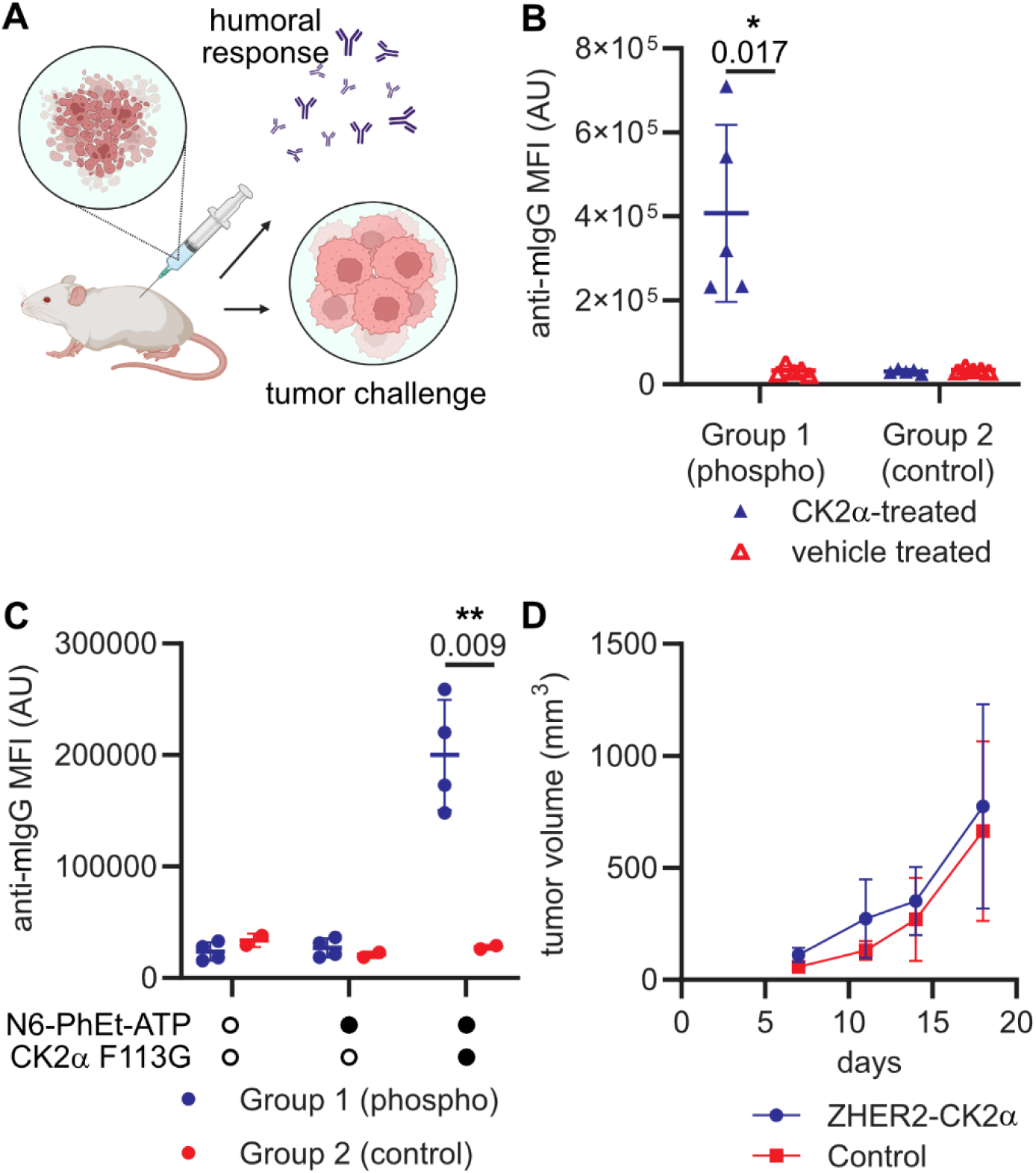
Whole-cell vaccination with hyperphosphorylated cell surfaces elicits a humoral but not a tumor-killing immune response. **(A)** Mice were immunized weekly for four weeks with heat-killed EMT-6 cells that had either been hyperphosphorylated with cell-tethered CK2α *in vitro* or treated with vehicle. Serum was collected and measured for binding to syngenic tumor cells or mice were challenged with live tumor cell implantation. **(B)** Binding of sera from five mice immunized with hyperphosphorylated EMT-6 cells (Group 1) or control cells (Group 2) to live EMT-6 pretreated with cell-tethered wild-type (CK2α -treated) or loss-of-function (control treated) CK2α *in vitro* shows significant increase in antibodies for cells treated with wild-type CK2α. Binding of mouse IgG was visualized by the binding of a rabbit anti-mIgG AlexaFluor647 conjugate. **(C)** Binding of immunized sera from Group 1 or Group 2 to EMT-6 cells pretreated with N6-phenylethyl ATP or vehicle and CK2α F113G or vehicle. Statistics for **(B)** and **(C)** were determined by corrected multiple t-tests. **(D)** Mice were then challenged with live EMT-6 cells that were either hyperphosphorylated with cell-tethered CK2α or treated with vehicle *in vitro* prior to implantation. Averaged tumor growth curves from mice immunized with hyperphosphorylated EMT-6 cells. For (**D**) error bars represent standard deviation of 5 mice per arm. * = corrected p value < 0.05. ** = corrected p value < 0.01. Corrected p-values are displayed above relevant comparisons. Non-significant comparisons are not displayed.

Given the robust antibody response generated by the mice, we wondered if the robust anti-phosphoprotein immune response may be sufficient to eradicate a tumor *in vivo*. Therefore, we challenged Group 1 immunized mice with hHER2+ EMT-6 cells pretreated with cell-tethered CK2α or vehicle (**Fig. 4D**). We did not observe any differences in tumor growth rate between the two treatments. Furthermore, we did not observe any differences in tumor growth rate for mice immunized with untreated wild-type EMT-6 cells, used for this immunization but not prior immunizations, (Group 2) or mock-immunized mice (Group 3) (**Fig. S11**). Collectively, these results suggest that cell surface CK2-substrates can be immunogenic, but that the immunity elicited is insufficient to mount an antitumoral response in this syngeneic model.

Finally, we sought to characterize the immune responses we observed. Given the robust degree of serum reactivity, we first proposed to identify the surface antigens on EMT-6 cells. We treated hHER2+ EMT-6 cells with cell-tethered CK2α, then bound sera from Group 1 or Group 2 animals to the surface, gently lysed the cells under non-denaturing conditions, and precipitated Ig-bound proteins using a Protein A/G magnetic bead. We eluted the digested bound proteins with trypsin and identified tryptic peptides by LC-MS (**Fig. 5A, Table S7**). Using this strategy, we confidently identified 217 unique proteins (≥ 2 unique peptides, ≤1% FDR) from Group 1, and only 2 unique proteins from Group 2 sera (**Fig. 5B**). Notably, CD44 was among the unique proteins identified from Group 1 sera. We observed that sera from multiple Group 1 mice but not Group 2 bound to *in vitro* phosphorylated recombinant murine (rm)CD44 (pCD44) by ELISA (**Fig. 5C**), and that Group 1 mice had increased reactivity towards pCD44 over rmCD44.

**Figure 5.**
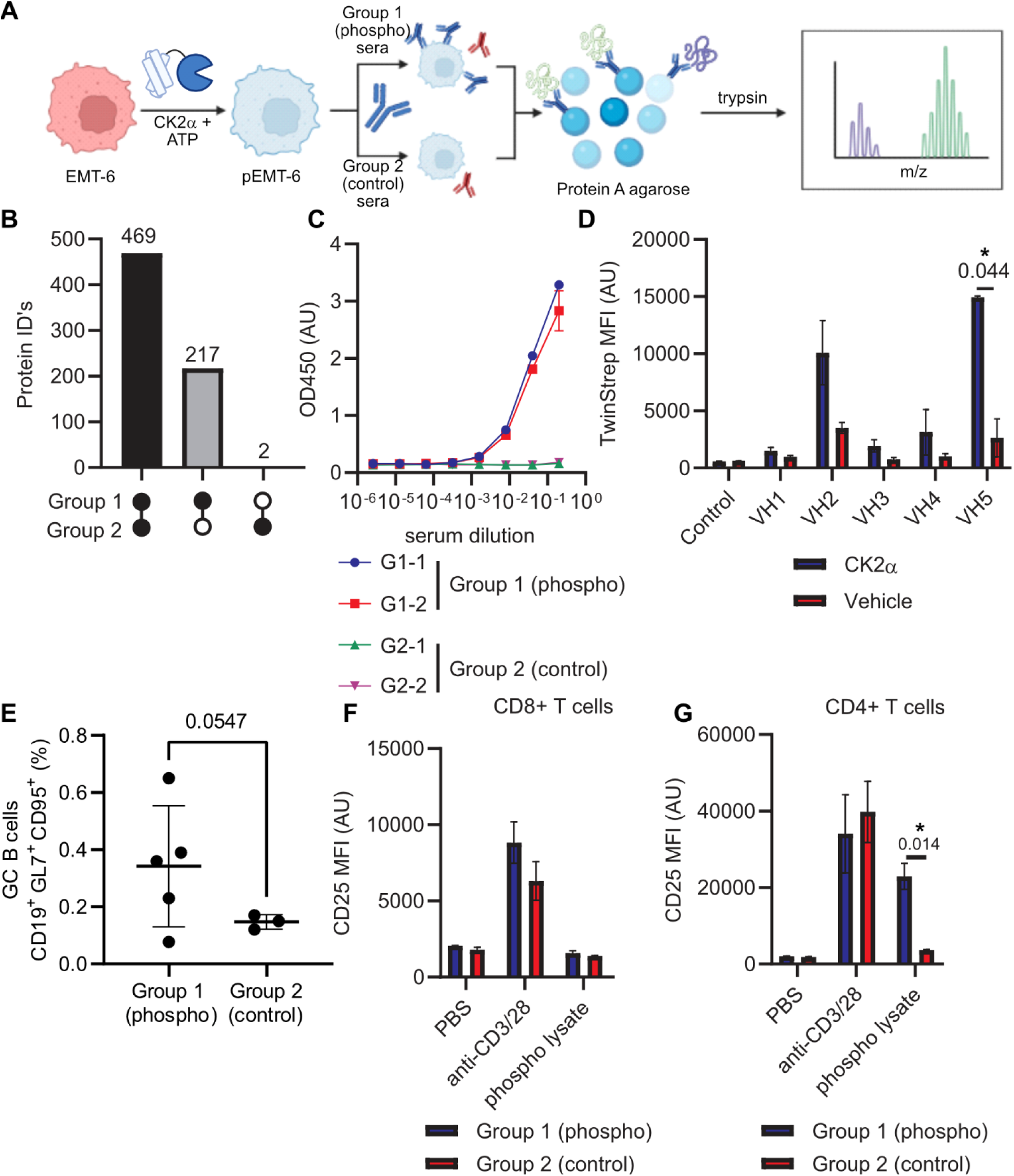
Immunization with CK2 surface substrates elicits a humoral immune response**. (A-D)** Immunogens were identified by binding sera from either Group 1 (wild-type ZHER2-CK2α) or Group 2 (inactive ZHER2-CK2α) mice to hHER2+ EMT-6 pretreated with cell-tethered CK2α, gently lysing the cells, precipitating mIgG on immobilized Protein A, and analyzing tryptic digests by LC-MS. **(B)** Identification of unique proteins from -immunized sera shows many more for the wild-type ZHER2-CK2α treated mice. **(C)** rmCD44 was phosphorylated *in vitro* with CK2 and tested for binding sera by ELISA shows positive sera for the wild-type ZHER2-CK2α treated mice. **(D)** VH genes were subcloned from splenocyte mRNA and ported into a scFv-based yeast-display vector for enrichment of phospho-CD44 binding clones. Top 5 enriched clones were expressed with a TwinStrep affinity tag and binding to hHER2+ EMT-6 cells pretreated with cell-tethered CK2α was quantified using a fluorescent Streptactin-XT. Statistics were determined by multiple t-tests corrected for multiple comparisons by the Holm-Sidak method. **(E-G)** Splenocytes from immunized mice were characterized by flow cytometry. **(E)** Germinal B cells were quantitated as a proportion of total B cells. Statistics determined by Student’s T test. **(F,G)** Splenocytes were stimulated with either antibody cocktails or hHER2+ EMT-6 cell, pretreated with cell-tethered CK2α, lysates and allowed to activate for 72 hours. Analysis of the activation state of CD8+ T cells **(F)** and CD4+ T cells **(G)** was measured by upregulation of CD25 by flow cytometry. Statistics for **(F,G)** were determined by corrected multiple T tests. * = corrected p value < 0.05. Corrected p-values are displayed above relevant comparisons. Non-significant comparisons are not displayed.

To identify specific pCD44-reactive sequences, we amplified the VH genes from mRNA isolated from the pooled splenocytes of Group 1 mice. We then subcloned the pooled population into a common light chain yeast display scFv library. We sorted the library with pCD44 by successive rounds of magnetic- and fluorescence-activated cell sorting to identify a subpopulation of pCD44-specific clones (**Fig. S12**). We sequenced this pooled population and cloned the top five most abundant scFv sequences. We found that two clones bound hHER2+ EMT-6 cells treated with cell-tethered CK2α, and that one was highly specific for CK2-mediated phosphorylation (**Fig. 5D, Fig. S13**).

We next characterized the cellular component of the immune response using splenocytes isolated from immunized animals. We quantified the number of germinal center B cells and measured their binding to rmCD44 and pCD44 simultaneously by flow cytometry (**Fig. 5E, Fig. S14**). We saw a modest increase (p = 0.054, Cohen’s d = 1.34) in the proportional population of GC B cells in Group 1 versus Group 2 animals, but observed that most CD44-reactive clones bound both rmCD44 and pCD44. We also probed the reactivity of splenic T cells in a mixed leukocyte reaction.^48^ We fed bulk splenocytes the lysates of heat-killed hHER2+ EMT-6, which had been pretreated with cell-tethered CK2α,and measured CD25 upregulation as a proxy for T cell activation upon binding splenic antigen presenting cells that took up the lysates (**Fig. 5F-G, Fig. S15**). We observed that CD4+ T cells upregulated CD25 in response to the surface phosphorylated lysates but that CD8+ T cells did not which may account for the strong antibody response but lack of antitumor response.

## Discussion

The tethering CK2α to the surface provides another example of how surface-tethering of enzymes increases their efficacy as has been shown with ligases,^34,35^ proteases,^36^ and sialidases,^37^ among other enzymes.^49^ This enzyme tethering strategy was very useful to investigate and enhance the substrate scope of this kinase. We identified several tumor-associated antigens that are surface substrates of CK2α, including CD44, Galectin-3, and CD109. Given that tumor cells are known to express CK2, we hypothesize that endogenous CK2 may hyperphosphorylate these proteins in tumors, which possess chronically high extracellular ATP, and thus provide therapeutically relevant phosphoantigens. We anticipate that the surface phosphoproteome will provide fertile ground for new tumor biomarkers, and that the tools and workflows devised here will help identify these biomarkers.

It was remarkable that these ecto-phosphoproteins elicited a humoral immune response and activation of CD4+ T cells without activating a CD8+ T cell response *in vivo*. We speculate that this is due to the fact that CD8+ T cells largely rely on engaging MHC-I peptides, which predominantly display intracellular proteins, and that only a select subset of MHC-I proteins effectively display these ecto-derived phosphopeptides. These factors likely limit the display of the phosphopeptides generated by ectokinase activities, and thus limit the CD8+ T cell response. However, we found the degree of phosphoprotein serum reactivity remarkable, including the phosphorylation-independent reactivity, suggesting antigenic spread did occur.

In summary, these studies collectively present both new methodologies for probing the extracellular phosphoproteome as well as evidence that extracellular phosphoproteins can be immunogenic. Due to the high intra-tumoral concentrations of ATP, it is reasonable to presume that ectokinases will be chronically active in that environment, resulting in a dramatic increase in specific phosphoproteins compared to the healthy tissue. We anticipate the chemoproteomic methods described here can uncover more tumor-associated phosphoprotein antigens that may eventually be leveraged therapeutically.

## Acknowledgements

The authors thank Kevin Leung and Jamie Byrnes for their assistance with proteomics discussions, and the Wells Lab broadly for helpful discussions and expertise. J.A.W. is supported by generous grants from NIH NCI 1P41CA196276, CA191018, and NIH GM097316 and commercial funding from Bristol Myers Squibb. C.S.D. is an AP Giannini Foundation Postdoctoral Fellow. L.L.K. was supported by the NIH F31 Ruth L. Kirschstein National Research Service Award (1F31CA247527). Data for this study were acquired at the Center for Advanced Light Microscopy at UCSF on CREST/C2 Confocal obtained using grants from the UCSF Program for Breakthrough Biomedical Research funded in part by the Sandler Foundation and the UCSF Research Resource Fund Award. Animal studies were supported by the HDFCCC Shared Resource Facility, Preclinical Therapeutics Core, through NIH (P30CA082103).

## RESOURCE AVAILABILITY

### Lead contact

- Further information and requests for resources and reagents should be directed to and will be fulfilled by the lead contact, Jim Wells.

### Materials availability

- The coding sequences for plasmids generated in this study are available in the supporting information for public use.

### Data and code availability

- Mass spectrometry-based proteomics data have been deposited via PRIDE^50^ with the identifier PXD050801 and are publicly available as of the date of publication. Accession numbers are listed in the key resources table. Full Western blots and representative flow cytometry data are available in the supplementary information. Microscopy images, raw Western blots, and raw flow cytometry data are available upon request.
- This paper does not report original code.
- Any additional information required to reanalyze the data reported in this paper is available from the lead contact upon request.

## EXPERIMENTAL MODEL

### Animals

All mice were 6–9-week-old female BALB/c mice obtained from Jackson Laboratories (stock# 00651). All work performed on mice was approved under the Institutional Animal Care and Use Committee (IACUC) protocol AN194778 at the University of California, San Francisco. Mice were housed with *ad libitum* food and water on a 12-hour light cycle at the UCSF Preclinical Therapeutics Core vivarium.

### Primary cell cultures

Splenocytes were isolated from the spleens of mice used in this study. Primary splenocytes were stored 20% fetal bovine serum (FBS) in RPMI-1640 in cryovials suspended in the vapor phase of liquid nitrogen. Splenocytes were cultured in RPMI-1640 supplemented with 10% FBS, 50 μM β-mercaptoethanol, and 10 U/mL recombinant IL-2 (R&D Systems, 402-ML-020/CF). Cells were cultured in a humidified incubator at 37 °C and 5% CO_2_.

### Cell lines

EMT-6 (ATCC, CRL-2755) cells are a murine mammary carcinoma derived from BALB/cRgl mice. EMT-6 hHER2+ cells are a human HER2-overexpressing derivative of the parental EMT-6 line and were generated in house using pHAGE-ERBB2 (Addgene, 116734). EMT-6 cells were maintained in adherent culture in DMEM supplemented with 10% FBS. SK-BR-3 cells are human mammary gland adenocarcinoma. SK-BR-3 (ATCC, HTB-30) cells were maintained in adherent culture in McCoy’s 5a medium supplemented with 10% FBS. Cells were cultured in a humidified incubator at 37 °C and 5% CO_2_.

ExpiCHO cells were cultured in ExpiCHO Expression medium, per the manufacturer’s instructions, in suspension culture in an 8% CO_2_ incubator on an orbital rotating platform at 120 rpm.

#### Microbe strains

EBY100 yeast (ATCC, MYA-4941) cells were cultured in YPD medium. BL21(DE3) *E. coli* (NEB, C2527H) were used for recombinant protein expression. A C43 cell derivative, C43(DE3) Pro+ Tum+, has been previously developed by our lab and described.^51^

## METHOD DETAILS

### Plasmid construction and cloning

Recombinant DNA for an optimized sequence of CK2α and ZHER2 was ordered from Integrated DNA Technologies (IDT) and is listed in **Table S9**. Gene fragments were subcloned into a pMAL-c6T (NEB, N0378S) expression vector by Gibson Assembly.

Constructs bearing a C-terminal cMyc tag were cloned by site-directed mutagenesis (NEB, M0554S) using oligonucleotides as primers bearing a 5’-overhang encoding the inserted sequence. Oligonucleotide sequences are listed in **Table S8**. Mutants of CK2α were cloned by site-direction mutagenesis using oligonucleotides as primers bearing a 5’-overhang encoding the mutagenic sequence. Truncation mutants were cloned likewise.

For plasmids encoding CK2α as a knob-in-hole Fc fusion, a fragment of CK2α recombinant DNA encoding residues 1 to 336 was subcloned into an in-house knob-and-hole pFUSE-based vector (Invivogen, pfuse-hg1fc1) with a Gly/Ser linker in between. Full sequence information is listed in **Table S9**.

For plasmids encoding CD44 expression constructs, a fragment of recombinant DNA encoding either residues 21 to 220 of human CD44 or 25 to 224 of murine CD44 were subcloned into an in-house Fc pFUSE-based vector (Invivogen, pfuse-hg1fc1) with a Gly/Ser linker in between. Full sequence information is listed in **Table S9**.

### Bacterial protein expression

Recombinant CK2α fusion proteins were expressed as maltose binding protein (MBP) conjugates in BL21(DE3) *E. coli* (NEB, C2527H) via autoinduction. Bacteria were harvested 18-24 hours after inoculation by centrifugation (4000 rcf, 20 min), lysed with B-PER Complete Protein Extraction Reagent (Thermo Scientific, 89821), and purified on amylose resin (NEB, E8021S) according to the manufacturer’s instructions.

Purified MBP-fusion proteins were treated with TEV-protease (NEB, P8112S) for 1 hour at room temperature, followed by removal of protease and cleaved-MBP using Ni-NTA resin (Gold Bio, H-350-5). Cleavage was validated by SDS-PAGE and size exclusion chromatography on a Agilent 1260 Infinity II system using an Agilent AdvanceBio SEC 300 Å 2.7 μm 4.6x300 mm column (Agilent Technologies, PL1580-5301) using the manufacturer’s recommended conditions. Excess maltose was removed from the purified protein by serial buffer exchange in 10 kDa MWCO spin columns. Purified proteins were stored at -80 °C in 10% glycerol in PBS.

Single-chain variable fragments were expressed C43(DE3) Pro+Tum+ *E. coli* (in house) via autoinduction. Bacteria were harvested 18-24 hours after inoculation by centrifugation (4000 rcf, 20 min), lysed with B-PER Complete Protein Extraction Reagent (Thermo Scientific, 89821), and purified on StrepTactin-XT resin (IBA Lifesciences, 2-5010-002) according to the manufacturer’s instructions. Excess biotin was removed from the purified protein by serial buffer exchange in 10 kDa MWCO spin columns. Purified proteins were stored at -80 °C in 10% glycerol in PBS.

### Mammalian protein expression

Fc fusion proteins were expressed in ExpiCHO cells using the ExpiCHO expression system kit (Thermo Scientific, A29133) according to the manufacturer’s instructions. Supernatants were collected by centrifugation (4000 rcf, 20 min). For CK2α-Fc knob constructs, proteins purified on Ni-excel resin (Cytiva, 17-3712-01) to enrich for a C-terminal His tag on the CK2α-Fc knob fusion protein using the manufacturer’s recommended conditions. Imidazole was removed from the purified protein by serial buffer exchange in 10 kDa MWCO spin columns (Millipore Sigma, UFC801096). For CD44-Fc constructs, proteins were purified on HiTrap Protein A cartridges (GE Lifesciences, 17-0402-01) using a peristaltic pump using the manufacturer’s recommended conditions. Eluates were neutralized by the addition of 1 M Tris base and buffer exchanged into PBS using 10 kDa MWCO spin columns (Millipore Sigma, UFC801096). Purified proteins were stored at -80 °C in 10% glycerol in PBS.

### ADP-Glo kinase activity assay

Kinase activity was measured in a generic kinase reaction buffer (20 mM Tris pH 7.4, 50 mM KCl, 10 mM MgCl2) using the ADP-Glo assay kit (Promega, V6930) and on a commercial CK2 substrate peptide (EMD Millipore, 12-330) under the manufacturer’s recommended conditions. Luminescence was read on a Tecan Infinite M200 Pro plate reader.

### Biolayer interferometry

Binding kinetics of ZHER2-CK2α were measured by biolayer interferometry (BLI) on an OctetRed384 using in-house made HER2-Fc-AviTag immobilized on streptavidin biosensors (Sartorius, 18-5021). Kinetic measurements were performed in PBS with 0.1% bovine serum albumin and 0.05% Tween 20.

### On-cell tethering of CK2**α** by flow cytometry

Cells were lifted from adherent monolayer culture with Versene (Fisher Scientific, 15-040-066) at 37 °C, harvested by centrifugation (500 rcf), and resuspended in PBS with 10 mM MgCl_2_ and 1 mM ATP (Sigma Aldrich, A1852-1VL) in the presence of cMyc-tagged CK2α fusion protein (1 μM). Cells were incubated at 37 °C for 30 min, washed three times with cold 1% BSA in PBS (500 rcf, 5 min), and incubated on ice for 30 min with anti-cMyc clone 9E10 AlexaFluor 647 conjugate (Thermo Fisher Scientific, MA1980A647) at a 1:100 dilution in 1% BSA in PBS. Cells were washed two times with cold 1% BSA in PBS (500 rcf, 5 min), once with PBS (500 rcf, 5 min), and incubated for 15 min in PBS with Propidium Iodide Ready Flow (Thermo Scientific, R37169) live/dead cell stain. Cells were analyzed on a Beckman Coulter CytoFLEX flow cytometer.

### On-cell tethering of CK2**α** by confocal microscopy

Approximately 5,000 EMT-6 hHER2+ cells were seeded in Ibidi 8-well microchamber glass slides (Ibidi, 80826) 24 hours prior to treatment and grown under standard conditions. On the day of treatment, adhered cells were washed with PBS and treated with cMyc-tagged CK2α or ZHER2-CK2α (1 μM) in PBS with 10 mM MgCl_2_ and 1 mM ATP (Sigma Aldrich, A1852-1VL). Cells were incubated at 37 °C for 30 min, washed three times with cold 1% BSA in PBS, and fixed in 10% neutral formalin for 20 minutes at room temperature in the dark. The cells were washed three times with PBS and then permeabilized with 0.1% Triton-X100 in PBS for 20 minutes at room temperature. Fixed and permeabilized cells were then washed three times with PBS and blocked with 10% heat-inactivated human serum for 1 hour at room temperature. Cells were then co-stained with anti-cMyc clone 9E10 AlexaFluor 647 conjugate (Thermo Fisher Scientific, MA1980A647) at a 1:50 dilution and anti-HER2 (trastuzumab biosimilar) AlexaFluor488 conjugate at 1:50 dilution in 1% BSA in PBS overnight at 4 °C. Cells were then washed three times with PBS, stained with 1 μg/mL DAPI (Cell Signaling Technologies, 4083S) in PBS for 5 min, and then washed three times with PBS before being immersed in Ibidi mounting medium (Ibidi, 50001). Cells were imaged on a Nikon Ti2 inverted fluorescence microscope with a Nikon C2 laser scanning confocal with 4 laser lines in the Center for Advanced Light Microscopy at the University of California, San Francisco.

### Activity of CK2**α** on purified proteins

Recombinant or purified proteins (1 mg/mL) were incubated in the presence of CK2α, or mutant, in PBS with 10 mM MgCl_2_ and either 1 mM ATP (Sigma Aldrich, A1852-1VL) or 1 mM ATP-γ-thiophosphate (Sigma Aldrich, A1388-25MG) for 30 min at 37 °C. Reactions were alkylated with 2 mM p-nitrobenzylmesylate (Abcam, ab138910) for 2 hours at room temperature, separated by SDS-PAGE, transferred to PVDF membranes, and blots were blocked with 5% BSA in TBS. For experiments using ATP, blots were probed with anti-phospho-CK2 substrate (pS/pT)DXE (Cell Signaling Technologies, 8738S). For experiments using ATP-γ-thiophosphate, blots were probed with anti-thiophosphate clone 51-8 (Abcam, ab92570) (1:15000), washed, probed with goat anti-rabbit IgG IRDye 800CW conjugate (LiCOR, 926-32211) (1:10000) and goat anti-mouse IgG 680RD conjugate (LiCOR, 926-68070) (1:10000). Blots were imaged on a LiCOR CLx.

### On-cell activity of CK2**α** and ZHER2-CK2**α** fusion proteins

EMT-6 hHER2+ or SK-BR-3 cells were harvested with Versene and harvested by centrifugation (500 rcf) and resuspended in PBS with 10 mM MgCl_2_ and 1 mM ATP-γ-thiophosphate (Sigma Aldrich, A1388-25MG) for 30 min at 37 °C in the presence of CK2α fusion proteins (1 μM) or vehicle. Cells were then washed three times with PBS and lysed in RIPA supplemented with HALT protease and phosphatase inhibitor (Thermo Fisher Scientific, 78440) and benzonase nuclease (Sigma Aldrich, E1014-25KU) for 30 min at 4 °C. Lysates were clarified by centrifugation (21000 rcf, 15 min) and protein concentration was quantitated by RapidGold BCA (Thermo Scientific, A53225). For experiments using ATP-γ-thiophosphate, normalized concentrations of lysates alkylated with 2 mM p-nitrobenzylmesylate (Abcam, ab138910) for 2 hours at room temperature. Normalized quantities of lysate were separated by SDS-PAGE and transferred to PVDF membranes. Blots were blocked with 5% BSA in TBS, probed with anti-thiophosphate clone 51-8 (Abcam, ab92570) (1:15000) and mouse anti-tubulin clone DM1A (Sigma Aldrich, T9026-.2ML) (1:5000), washed, probed with goat anti-rabbit IgG IRDye 800CW conjugate (LiCOR, 926-32211) (1:10000) and goat anti-mouse IgG 680RD conjugate (LiCOR, 926-68070) (1:10000). Blots were imaged on a LiCOR CLx.

### Ecto-phosphoproteomics enrichment

EMT-6 hHER2+ or SK-BR-3 cells were harvested with Versene and harvested by centrifugation (500 rcf) and resuspended in PBS with 10 mM MgCl_2_ and 1 mM ATP-γ-thiophosphate (Sigma Aldrich, A1388-25MG) for 30 min at 37 °C. For EMT-6 hHER2+ experiments, cells were incubated in the presence of cell-tethered ZHER2-CK2α or a D156A analogue (1 μM). For SK-BR-3 experiments, cells were cultured in the presence of CX-4945 (Biosynth International, FC30749) (5 μM) or an equal volume of DMSO. Cells were then washed three times with PBS and lysed in RIPA supplemented with HALT protease and phosphatase inhibitor (Thermo Fisher Scientific, 78440) and benzonase nuclease (Sigma Aldrich, E1014-25KU) for 30 min at 4 °C. Lysates were clarified by centrifugation (21000 rcf, 15 min) and protein concentration was quantitated by RapidGold BCA (Thermo Scientific, A53225). Normalized quantities of lysate, typically 400 μg in 400 μL, were immobilized on 100 μL Ultralink Iodoacetyl Resin (Thermo Scientific, 53155B) prewashed with RIPA at room temperature for 16 hours.

Resin was washed on a vacuum manifold in 2 mL columns, being careful to not let the resin dry, with three successive washes of 1 mL each of RIPA, PBS with 1 M NaCl, and 50 mM ammonium bicarbonate with 2 M urea. Resins were then alkylated and digested on-bead using Preomics iST 96x kits (Preomics, P.O.00027) according to the manufacturer’s protocol and eluted peptides were dried in vacuo. Peptides were resuspended in running solvent and quantitated using a Pierce Colorimetric Peptide Quantitation Assay (Thermo Fisher Scientific, 23275).

### Liquid chromatography mass spectrometry

Desalted peptides (200 ng) were loaded onto a timsTOF Pro equipped with a CaptiveSpray source and a nanoElute line (Bruker). The peptides were separated on a ReproSil C18 1.5 μM 100 Å 250 mm column (PepSep, PSC-25-150-15-UHP-nc) using a stepwise linear gradient method with H2O in 0.1% Formic acid and acetonitrile with 0.1% formic acid (solvent B): 5-30% solvent B for 90 min at 0.5 μl/min, 30-35% solvent B for 10 min at 0.6 μl/min, 35-95% solvent B for 4 min at 0.5 μl/min, 95% hold for 4 min at 0.5 μl/min). Acquired data was collected in a data-dependent acquisition mode with ion mobility activated in PASEF mode. MS and MS/MS spectra were collected with m/z ranging from 100 to 1700 in positive mode.

### LC-MS data analysis

All acquired data was searched using PEAKS online Xpro 1.6 (Bioinformatics Solutions Inc.; Ontario, Canada).^52^ Spectral searches were performed using a custom FASTA-formatted dataset containing either a human or murine version of the Swissprot-reviewed species proteome file with gene ontology localized the plasma membrane (downloaded from Uniprot knowledge database, #entries). A precursor mass error tolerance was set to 20 ppm and a fragment mass error tolerance was set at 0.03 ppm. Peptides, ranging from 6 to 45 amino acids in length, were searched in semi-specific tryptic digest mode with a maximum of two missed cleavages. Carbidomethylation (+57.02 Da) on cysteines was set as a static modification and both methionine oxidation (+15.99 Da) and N-terminal acetylation (+42.02 Da) were set as a variable modification. Lastly, peptides were filtered based on a false discovery rate (FDR) of 1%.

### Immunoprecipitation of CD44

EMT-6 hHER2+ cells were harvested with Versene and harvested by centrifugation (500 rcf) and resuspended in PBS with 10 mM MgCl_2_ and 1 mM ATP-γ-thiophosphate (Sigma Aldrich, A1388-25MG) for 30 min at 37 °C in the presence of cell-tethered ZHER2-CK2α or a D156A analogue (1 μM). Cells were then washed three times with PBS and lysed in 1% NP-40 in TBS supplemented with HALT protease and phosphatase inhibitor (Thermo Fisher Scientific, 78440) and benzonase nuclease (Sigma Aldrich, E1014-25KU) for 30 min at 4 °C. Lysates were clarified by centrifugation (21000 rcf, 15 min) and protein concentration was quantitated by RapidGold BCA (Thermo Scientific, A53225). Concentrations of lysates were normalized, alkylated with 2 mM p-nitrobenzylmesylate (Abcam, ab138910) for 1 hour at room temperature, and 200 µg of lysate was immunoprecipitated with 40 µL Protein A dynabeads (Fisher Scientific, 10-001-D) pre-coupled to 8 µg anti-CD44 (Fisher Scientific, 15675-1-AP) with BS3 crosslinker (Thermo Scientific, A39266). Lysates were immunoprecipitated overnight, washed three times with 1% NP-40 in TBS, and eluted with SDS buffer. Immunoprecipitates were separated by SDS-PAGE in duplicate and transferred to PVDF membranes. Blots were blocked with 5% BSA in TBS, probed with anti-thiophosphate clone 51-8 (Abcam, ab92570) (1:15000) or anti-CD44 (Fisher Scientific, 15675-1-AP) (1:500), washed, probed with goat anti-rabbit IgG IRDye 800CW conjugate (LiCOR, 926-32211) (1:10000). Blots were imaged on a LiCOR CLx.

### Phosphosite identification on purified phosphoproteins

Either human or murine CD44-Fc-AviTag N-terminal domain protein (1 mg/mL) was incubated in the presence of CK2α, or mutant, in PBS with 10 mM MgCl_2_ and 1 mM ATP (Sigma Aldrich, A1852-1VL) for 30 min at 37 °C. CD44 was immobilized on High Capacity Neutravidin Agarose (Thermo Fisher Scientific, 29204) by incubating reactions on agarose for 1 hour at room temperature. Resin was washed on a vacuum manifold in 2 mL columns, being careful to not let the resin dry, with three successive washes of 1 mL each of RIPA, PBS with 1 M NaCl, and 50 mM ammonium bicarbonate with 2 M urea. Resins were then alkylated and digested on-bead using Preomics iST 96x kits (Preomics, P.O.00027) according to the manufacturer’s protocol and eluted peptides were dried in vacuo. Peptides were resuspended in running solvent and quantitated using a Pierce Colorimetric Peptide Quantitation Assay (Thermo Fisher Scientific, 23275). Peptides were separated and analyzed as described above, with the addition that phosphorylation (+79.97 Da) was included as a variable modification and the spectra were searched against a FASTA-formatted dataset containing the sequences of **the purified proteins used.**

### Mouse immunization experiments

Wild-type or hHER2+ EMT-6 cells were harvested with Versene and harvested by centrifugation (500 rcf) and resuspended in PBS with 10 mM MgCl_2_ and 1 mM ATP (Sigma Aldrich, A1852-1VL) for 30 min at 37 °C in the presence of cell-tethered ZHER2-CK2α or a D156A analogue (1 μM). Cells were then washed three times with PBS and then resuspended at 1e7 cell per mL in PBS and killed by heat-shock at 47 °C for 1 hour with shaking at 900 rpm.^45^ Heat-killed lysates were immediately flash-frozen until use.

BALB/c mice received subcutaneous injections of 100 µL heat-killed lysate in alternating flanks weekly for four weeks. At the end of the four weeks, mice were euthanized according to standard euthanasia protocols per IACUC protocol AN194778at the University of California, San Francisco. Blood and spleens were harvested for later use.

Serum was separated from whole blood by centrifugation. Serum was then aliquoted and flash-frozen in liquid nitrogen. Frozen aliquots were stored at -80 °C.

Splenocytes were harvested by homogenizing spleens using a 45 μm nylon mesh filter and the rubber end of a 3 mL syringe plunger in 2 mL PBS. The dissociated cells were harvested by centrifugation (600 rcf, 5 min). Pellets were resuspended in 2 mL ACK lysis buffer (Fisher Scientific, A1049201) and incubated at room temperature for 3 min, at which point 10 mL PBS was added to neutralize the lysis. Cells were pelleted by centrifugation (600 rcf, 5 min), washed with 10 mL PBS again, and resuspended in 1 mL cryopreservation medium (Biolife, 210102) for gradual cooling and long-term storage in liquid nitrogen vapor phase.

### Serum flow cytometry

EMT-6 hHER2+ cells were harvested with Versene and harvested by centrifugation (500 rcf) and resuspended in PBS with 10 mM MgCl_2_ and either 1 mM ATP (Sigma Aldrich, A1852-1VL) or 1 mM N6-phenylethyl-ATP (Biolog, P 012-05) for 30 min at 37°C in the presence of CK2α fusion proteins (1 μM) or vehicle. Cells were washed three times with cold 1% BSA in PBS (500 rcf, 5 min) and incubated on ice for 30 min with diluted serum (1:50) from immunized animals. Cells were washed three times with cold 1% BSA in PBS (500 rcf, 5 min) and incubated on ice for 30 min with goat anti-mouse IgG AlexaFluor 647 conjugate (Thermo Fisher Scientific, A28181) at a 1:200 dilution in 1% BSA in PBS. Cells were washed two times with cold 1% BSA in PBS (500 rcf, 5 min), once with PBS (500 rcf, 5 min), and incubated for 15 min in PBS with Propidium Iodide Ready Flow (Thermo Scientific, R37169) live/dead cell stain. Cells were analyzed on a Beckman Coulter CytoFLEX flow cytometer.

### Mouse immunization then challenge experiments

Mice were immunized as before. EMT-6 hHER2+ cells were harvested with Versene and harvested by centrifugation (500 rcf) and resuspended in PBS with 10 mM MgCl_2_ and 1 mM ATP (Sigma Aldrich, A1852-1VL) for 30 min at 37 °C in the presence of cell-tethered ZHER2-CK2α or a D156A analogue (1 μM). Cells were then resuspended at 1e7 cell per mL in PBS and injected subcutaneously into the flanks of immunized or naïve BALB/c mice (1e6 cell/100ul in 1:1 PBS:Matrigel). Tumor growth was assessed biweekly by 2D caliper measurement and volume was calculated according to the volume of an ellipsoid ((width)^2^ x length x 0.52).

### Immunoprecipitation of antigen proteins

EMT-6 hHER2+ cells were harvested with Versene and harvested by centrifugation (500 rcf) and resuspended in PBS with 10 mM MgCl_2_ and either 1 mM ATP (Sigma Aldrich, A1852-1VL) 30 min at 37 °C in the presence of CK2α fusion proteins (1 μM) or vehicle. Cells were washed three times with cold 1% BSA in PBS (500 rcf, 5 min) and incubated on ice for 30 min with diluted serum (1:50) from immunized animals. Cells were then washed three times with PBS and lysed in 1% NP-40 in TBS supplemented with HALT protease and phosphatase inhibitor (Thermo Fisher Scientific, 78440) and benzonase nuclease (Sigma Aldrich, E1014-25KU) for 30 min at 4 °C. Lysates were clarified by centrifugation (21000 rcf, 15 min) and protein concentration was quantitated by RapidGold BCA (Thermo Scientific, A53225). 500 µg of lysate was immunoprecipitated with 40 µL Protein A dynabeads (Fisher Scientific, 10-001-D).

Lysates were immunoprecipitated overnight, washed three times with 1% NP-40 in TBS, and peptides were digested on-bead using Preomics iST 96x kits (Preomics, P.O.00027) according to the manufacturer’s protocol. Peptides were separated and analyzed as described above.

### Serum ELISA

A 384 well MaxiSorp plate (Thermo Fisher Scientific, 464718) was precoated with 50 µL of 0.5 µg/mL NeutrAvidin (Fisher Scientific, PI31000) and blocked with 1% BSA in PBS. Recombinant mCD44-Fc-AviTag (200 nM) in ELISA buffer (PBS with 0.2% BSA in 0.05% Tween-20) was immobilized on the plate for 30 min are room temperature. Residual NeutrAvidin binding sites were then blocked with 1 μM biotin in ELISA buffer.

The plate was washed three times, and either CK2α or a D156A analogue was added to each well (1 μM) in PBS with 10 mM MgCl_2_ and 1 mM ATP (Sigma Aldrich, A1852-1VL). Phosphorylation reactions were allowed to proceed for 30 min at 37 °C. The plate was washed three times and serum dilutions in ELISA buffer were added to each well in triplicate and allowed to bind for 1 hour at room temperature. The plate was washed three times and anti-mouse IgG-HRP (Thermo Fisher Scientific, 31430) (1:5000) in ELISA buffer and was incubated at room temperature for 30 min. The plate was washed three times and TMB (VWR International, 50-76-03) substrate was added. The TMB reaction was quenched by the addition of an equal volume of 1 M phosphoric acid. Absorbance was read at 450 nm using a Tecan Infinite M200Pro plate reader.

### Yeast display library generation

Splenocyte derived mRNA was harvested using a QIAmp RNA mini kit (Qiagen, 52304) and converted to cDNA using RNA to cDNA EcoDry Premix (Takara, 639547), both according to manufacturer’s instructions. A majority of murine VH genes were then amplified using a custom-mixed primer pool based on literature protocols.^53^ Adapter sequences for homology-directed recombination into an in-house yeast display vector, were added in a second PCR step. Oligonucleotide sequences can be found in **Table S8**.

Yeast were transformed by electroporation with PCR product and digested vector containing a common light chain based on the frequently used non-binding MOPC-21^54^ clone according to literature protocols.^55^ Transformants were cultured in SDCAA media.

### Yeast display sorting

Yeast were induced in SGCAA according to literature protocols.^55^ Yeast libraries were subjected to two successive rounds of magnetic activated cell sorting (MACS) followed by one round of fluorescence activated cell sorting (FACS). pCD44 was generated by pretreatment of recombinant murine CD44-Fc-AviTag with CK2α in PBS with 10 mM MgCl_2_ and 1 mM ATP (Sigma Aldrich, A1852-1VL). DNA was harvested from the libraries before and after sorting using Zymoprep Yeast Plasmid Miniprep II (Zymo Research, D2004). VH sequences were amplified by PCR and submitted to Genewiz for Amplicon NGS sequencing.

For MACS selections, 1e8 yeast were harvested by centrifugation (2000 rcf, 3 min) and resuspended in cold selection buffer (1% BSA and 0.02% Tween-20 in PBS). Yeast were incubated with either 100 nM pCD44 (positive) or 100 nM murine CD44-Fc (negative) for 30 min on ice, ensuring that copy numbers (estimating 50,000 scFv displayed per yeast) were at less than 1% potential ligand depletion. Cells were washed three times with selection buffer and labeled with 10 μL streptavidin MACS microbeads (Miltenyi Biotec, 130-048-102) (sufficient for 1e7 cells). Cells were washed three times with selection buffer and purified on LS-columns (Miltenyi Biotec) according to the manufacturer’s instructions for positive and negative selections.

For FACS selections, 5e6 yeast were harvested by centrifugation (2000 rcf, 3 min) resuspended in cold FACS buffer (3% BSA in PBS). Yeast were incubated with 50 nM pCD44 for 30 min on ice, ensuring that copy numbers (estimating 50,000 scFv displayed per yeast) were at less than 1% potential ligand depletion. Cells were washed three times with selection buffer and labeled with streptavidin AlexaFluor647 conjugate (Fisher Scientific, S21374) (1:200) and anti-cMyc clone 9E10 AlexaFluor 488 conjugate (Thermo Scientific, MA1-980-A488) (1:100) for 30 min on ice. Cells were washed three times with selection buffer and resuspended in selection buffer. Cells were sorted on a BD FACS Aria II, collecting the top 2% of expression and binders.

### Yeast display analysis by flow cytometry

Yeast libraries were harvested by centrifugation (2000 rcf, 3 min) resuspended in cold 1% BSA in PBS. Yeast were incubated with 50 nM pCD44, streptavidin AlexaFluor647 conjugate (Fisher Scientific, S21374) (1:200), and anti-cMyc clone 9E10 AlexaFluor 488 conjugate (Thermo Scientific, MA1-980-A488) (1:100) for 30 min on ice. Cells were washed three times with selection buffer and resuspended in PBS. Cells were analyzed on a Beckman Coulter CytoFLEX flow cytometer.

### On-cell binding of anti-pCD44 scFv’s

EMT-6 hHER2+ cells were harvested with Versene and harvested by centrifugation (500 rcf) and resuspended in PBS with 10 mM MgCl_2_ and 1 mM ATP (Sigma Aldrich, A1852-1VL) 30 min at 37 °C in the presence of CK2α fusion proteins (1 μM) or vehicle. Cells were washed three times with cold 1% BSA in PBS (500 rcf, 5 min) and incubated on ice for 30 min with 100 nM scFv and StrepTactin-XT DY647 conjugate (IBA Lifesciences, 2-1568-050) at a 1:200 dilution in 1% BSA in PBS. Cells were washed two times with cold 1% BSA in PBS (500 rcf, 5 min), once with PBS (500 rcf, 5 min), and incubated for 15 min in PBS with Propidium Iodide Ready Flow (Thermo Scientific, R37169) live/dead cell stain. Cells were analyzed on a Beckman Coulter CytoFLEX flow cytometer.

### Splenocyte quantitation

Splenocytes were thawed into 8 mL RPMI-1640 with 10% FBS, harvested by centrifugation (150 rcf, 8 min), and resuspended in complete growth media. Splenocytes were filtered through a 30 µm Nylon mesh filter into a 6 well dish and allowed to recover overnight. Splenocytes were then harvested by centrifugation, resuspended in Fc block in PBS for 10 min (BD Bioscience, 564220), and then in a GC B cell cocktail including: rat anti-mouse CD19 clone 1D3 PerCP-Cy5.5 conjugate (Fisher Scientific, BDB551001), hamster anti-mouse CD95 clone JO2 PE conjugate (Fisher Scientific, BDB554258), rat anti-mouse GL-7 Pacific Blue conjugate (BioLegend, 144614), pCD44-Fc-AviTag precomplexed with Streptavidin AlexaFluor 647 conjugate, and mCD44-Fc-AviTag precomplexed with Streptavidin AlexaFluor 488 conjugate. All antibodies were used at the manufacturer’s recommended dilutions. Splenocytes were stained for 30 min on ice, washed two times with cold 1% BSA in PBS (500 rcf, 5 min), once with PBS (500 rcf, 5 min), and resuspended in PBS for analysis. Cells were analyzed on a Beckman Coulter CytoFLEX flow cytometer compensated using single-color controls.

### Splenocyte activation

Splenocytes were thawed into 8 mL RPMI-1640 with 10% FBS, harvested by centrifugation (150 rcf, 8 min), and resuspended in complete growth media. Splenocytes were filtered through a 30 µm Nylon mesh filter into a 6 well dish and allowed to recover overnight. Splenocytes were then counter, harvested, and resuspended at 1e7 cells per mL. Aliquots of 100 μL were distributed in triplicate into a 96 well plate and stimulants were added in 10 μL PBS, being vehicle, 100 nM PMA (Millipore Sigma, P1585-1MG) plus 1 μg/mL ionomycin (Thermo Scientific, I24222), anti-murine CD3/28 T cell stimulatory dynabeads (Fisher Scientific, 11-456-D), or phosphorylated EMT-6 lysate (prepared as described in the Mouse Immunization protocols). Cells were incubated for 3 days, harvested by centrifugation, resuspended in Fc block in PBS for 10 min (BD Bioscience, 564220), and then in a T cell activation analysis cocktail including: anti-mouse CD3 clone 17A2 APC conjugate (BioLegend, 100235), rat anti-mouse CD4 AlexaFluor700 conjugate (BioLegend, 100429), anti-mouse CD8a clone 53-6.7 Pacific Blue conjugate (BioLegend, 480007), rat anti-mouse CD25 clone PC61 BV605 conjugate (BioLegend, 102035). All antibodies were used at the manufacturer’s recommended dilutions. Splenocytes were stained for 30 min on ice, washed two times with cold 1% BSA in PBS (500 rcf, 5 min), once with PBS (500 rcf, 5 min), and resuspended in PBS for analysis. Cells were analyzed on a Beckman Coulter CytoFLEX flow cytometer compensated using single-color controls.

## QUANTIFICATION AND STATISTICAL ANALYSES

- Statistical analyses were performed in GraphPad Prism 10 using ensuite analysis packages. The specific tests are detailed in the figure captions containing the comparison.

All flow cytometry data was analyzed at https://flowread.io

### KEY RESOURCES TABLES

| REAGENT or RESOURCE | SOURCE | IDENTIFIER |
| --- | --- | --- |
| Antibodies |  |  |
| anti-cMyc clone 9E10 AlexaFluor 647 | Thermo Fisher Scientific | MA1980A647 |
| anti-HER2 (trastuzumab biosimilar) AlexaFluor 488 | R&D Systems | FAB9589G-100UG |
| anti-thiophosphate ester clone 51-8 | Abcam | ab92570 |
| goat anti-rabbit IRDye 800CW conjugate | LiCOR | 926-32211 |
| goat anti-mouse 680RD conjugate | LiCOR | 926-68070 |
| mouse anti-tubulin clone DM1A | Sigma Aldrich | T9026-.2ML |
| anti-phospho-CK2 substrate (pS/pT)DXE | Cell Signaling Technologies | 8738S |
| anti-CD44, polyclonal | Thermo Scientific | 15675-1-AP |
| goat anti-mouse IgG AlexaFluor 647 conjugate | Thermo Fisher Scientific | A28181 |
| goat anti-mouse IgG HRP conjugate | Thermo Fisher Scientific | 31430 |
| anti-cMyc clone 9E10 AlexaFluor 488 conjugate | Thermo Scientific | MA1-980-A488 |
| rat anti-mouse CD19 clone 1D3 PerCP-Cy5.5 conjugate | Fisher Scientific | BDB551001 |
| hamster anti-mouse CD95 clone JO2 PE conjugate | Fisher Scientific | BDB554258 |
| rat anti-mouse GL-7 Pacific Blue conjugate | BioLegend | 144614 |
| anti-mouse CD3 clone 17A2 APC conjugate | BioLegend | 100235 |
| rat anti-mouse CD4 AlexaFluor700 conjugate | BioLegend | 100429 |
| anti-mouse CD8a clone 53-6.7 Pacific Blue conjugate | BioLegend | 480007 |
| rat anti-mouse CD25 clone PC61 BV605 conjugate | BioLegend | 102035 |
| Bacterial and virus strains |  |  |
| BL21(DE3) |  | C2527H |
| C43(DE3) Pro+Tum+ |  | Hornsby, et al.<sup>51</sup> |
| Chemicals, peptides, and recombinant proteins |  |  |
| rIL-2 | R&D Systems | 402-ML-020/CF |
| KLD enzyme mix | NEB | M0554S |
| B-PER Complete Protein Extraction Reagent | Thermo Scientific | 89821 |
| amylose resin | NEB | E8021S |
| TEV protease | NEB | P8112S |
| Ni-NTA agarose resin | GoldBio | H-350-5 |
| Ni-excel sepharose resin | Cytiva | 17-3712-01 |
| CK2 substrate peptide | EMD Millipore | 12-330 |
| Propidium Iodide Ready Flow Reagent | Thermo Scientific | R37169 |
| Adenosine triphosphate disodium salt, hydrate | Sigma Aldrich | A1852-1VL |
| DAPI | Cell Signaling Technologies | 4083S |
| Ibidi mounting media | Ibidi | 50001 |
| Casein, dephosphorylated | Sigma Aldrich | C4032-100MG |
| Mouse CD44[21-256] murine Fc fusion | R&D Systems | 6127-CD-050 |
| Mouse CD44[21-256] human Fc fusion | In house | This work |
| Human CD44[20-255] human Fc fusion | In house | This work |
| CD109 | R&D Systems | 4385-CD-050 |
| Galectin-3 | R&D Systems | 8259-GA-050 |
| p-nitrobenzylmesylate | Abcam | ab138910 |
| HALT protease and phosphatase inhibitor cocktail | Thermo Fisher Scientific | 78440 |
| Benzonase nuclease | Sigma-Aldrich | E1014-25KU |
| CX-4945 | Biosynth International | FC30749 |
| Ultralink Iodoacetyl Resin | Thermo Scientific | 53155B |
| HiTrap Protein A cartridges | GE Lifesciences | 17-0402-01 |
| High Capacity NeutrAvidin Agarose | Thermo Fisher Scientific | 29204 |
| ACK lysis buffer | Fisher Scientific | A1049201 |
| Cryopreservation medium | Biolife | 210102 |
| N6-phenylethyl-ATP | Biolog | P 012-05 |
| NeutrAvidin | Fisher Scientific | PI31000 |
| TMB Substrate | VWR International | 50-76-03 |
| StrepTactin XT 4Flow Resin | IBA Lifesciences | 2-5010-002 |
| Streptavidin Microbeads | Miltenyi Biotec | 130-048-102 |
| Streptavidin AlexaFluor 647 conjugate | Fisher Scientific | S21374 |
| StrepTactin XT DY647 | IBA Lifesciences | 2-1568-050 |
| Fc Block | BD Bioscience | 564220 |
| phorbol-12-myristate-13-acetate (PMA) | Millipore Sigma | P1585-1MG |
| ionomycin | Thermo Scientific | I24222 |
| Mouse T cell Activating CD3/CD28 dynabeads | Fisher Scientific | 11-456-D |
| Critical commercial assays |  |  |
| ADP-Glo Kinase Assay | Promega | V6930 |
| Streptavidin biosensors | Sartorius | 18-5021 |
| RapidGold BCA | Thermo Fisher | A53225 |
| Pierce Colorimetric Peptide Quantitation Assay | Thermo Fisher Scientific | 23275 |
| Preomics iST kit | Preomics | P.O.00027 |
| QIAmp RNA blood mini kit | Qiagen | 52304 |
| RNA to cDNA EcoDry Premix (Double Primed) | Takara | 639547 |
| Deposited data |  |  |
| Proteomics datasets |  |  |
| Experimental models: Cell lines |  |  |
| EMT-6 | ATCC | CRL-2755 |
| EMT-6 hHER2+ | in-house |  |
| SK-BR-3 | ATCC | HTB-30 |
| EBY100 | ATCC | MYA-4941 |
| Experimental models: Organisms/strains |  |  |
| Balb/c mice | UCSF animal |  |
| Oligonucleotides |  |  |
| See **Table S8** for oligonucleotide sequences |  |  |
| Recombinant DNA |  |  |
| pMAL-c6T | NEB | N0378S |
| CSNK21 | IDT | This work |
| pFUSE | Invivogen | pfuse-hg1fc1 |
| pHAGE-ERBB2 | Addgene | 116734 |
| mCD44 | IDT | This work |
| hCD44 | IDT | This work |
| pAAG263 | in-house | Cole's paper |
| See **Table S9** for full DNA sequences |  |  |
| Software and algorithms |  |  |
| PEAKS |  |  |
| Other |  |  |
| AdvanceBio SEC 300 Å 2.7 µm 4.6x300 mm column | Agilent Technologies | PL1580-5301 |
| ExpiCHO expression system kit | Thermo Scientific | A29133 |
| Ibidi 8-well microchamber glass slides | Ibidi | 80826 |
| ReproSil C18 1.5 µM 100 Å 250 mm column | PepSep | PSC-25-150-15-UHP-nc |

## Notes

### Competing Interest Statement

The authors have declared no competing interest.

## References

1. Cobbold, M., De La Peña, H., Norris, A., Polefrone, J.M., Qian, J., English, A.M., Cummings, K.L., Penny, S., Turner, J.E., Cottine, J., et al. (2013). MHC class I-associated phosphopeptides are the targets of memory-like immunity in leukemia. Sci. Transl. Med. 5, 1–11. 10.1126/scitranslmed.3006061.

2. Schumacher, T.N., and Schreiber, R.D. (2015). Neoantigens in cancer immunotherapy. Science (80-. ). 348, 69–74. 10.1126/science.aaa4971.

3. Malaker, S.A., Penny, S.A., Steadman, L.G., Myers, P.T., Loke, J.C., Raghavan, M., Bai, D.L., Shabanowitz, J., Hunt, D.F., and Cobbold, M. (2017). Identification of glycopeptides as posttranslationally modified neoantigens in Leukemia. Cancer Immunol. Res. 5, 376–384. 10.1158/2326-6066.CIR-16-0280.

4. Doyle, H.A., Zhou, J., Wolff, M.J., Harvey, B.P., Roman, R.M., Gee, R.J., Koski, R.A., and Mamula, M.J. (2006). Isoaspartyl post-translational modification triggers anti-tumor T and B lymphocyte immunity. J. Biol. Chem. 281, 32676–32683. 10.1074/jbc.M604847200.

5. Apostolopoulos, V., Yuriev, E., Ramsland, P.A., Halton, J., Osinski, C., Li, W., Plebanski, M., Paulsen, H., and McKenzie, I.F.C. (2003). A glycopeptide in complex with MHC class I uses the GalNAc residue as an anchor. Proc. Natl. Acad. Sci. U. S. A. 100, 15029–15034. 10.1073/pnas.2432220100.

6. Falzoni, S., Donvito, G., and Di Virgilio, F. (2013). Detecting adenosine triphosphate in the pericellular space. Interface Focus 3. 10.1098/rsfs.2012.0101.

7. Conley, J.M., Radhakrishnan, S., Valentino, S.A., and Tantama, M. (2017). Imaging extracellular ATP with a genetically-encoded, ratiometric fluorescent sensor. PLoS One 12, 1–24. 10.1371/journal.pone.0187481.

8. Pellegatti, P., Raffaghello, L., Bianchi, G., Piccardi, F., Pistoia, V., and Di Virgilio, F. (2008). Increased level of extracellular ATP at tumor sites: In vivo imaging with plasma membrane luciferase. PLoS One 3, 1–9. 10.1371/journal.pone.0002599.

9. Di Virgilio, F., Sarti, A.C., Falzoni, S., De Marchi, E., and Adinolfi, E. (2018). Extracellular ATP and P2 purinergic signalling in the tumour microenvironment. Nat. Rev. Cancer 18, 601–618. 10.1038/s41568-018-0037-0.

10. Hu, L.P., Zhang, X.X., Jiang, S.H., Tao, L.Y., Li, Q., Zhu, L.L., Yang, M.W., Huo, Y.M., Jiang, Y.S., Tian, G.A., et al. (2019). Targeting purinergic receptor P2Y2 prevents the growth of pancreatic ductal adenocarcinoma by inhibiting cancer cell glycolysis. Clin. Cancer Res. 25, 1318–1330. 10.1158/1078-0432.CCR-18-2297.

11. Gilbert, S., Oliphant, C., Hassan, S., Peille, A., Bronsert, P., Falzoni, S., Di Virgilio, F., McNulty, S., and Lara, R. (2019). ATP in the tumour microenvironment drives expression of nfP2X 7 , a key mediator of cancer cell survival. Oncogene 38, 194–208. 10.1038/s41388-018-0426-6.

12. Feng, L.L., Cai, Y.Q., Zhu, M.C., Xing, L.J., and Wang, X. (2020). The yin and yang functions of extracellular ATP and adenosine in tumor immunity. Cancer Cell Int. 20, 1–11. 10.1186/s12935-020-01195-x.

13. Bianchi, G., Vuerich, M., Pellegatti, P., Marimpietri, D., Emionite, L., Marigo, I., Bronte, V., Di Virgilio, F., Pistoia, V., and Raffaghello, L. (2014). ATP/P2X7 axis modulates myeloid-derived suppressor cell functions in neuroblastoma microenvironment. Cell Death Dis. 5, 1–12. 10.1038/cddis.2014.109.

14. De Marchi, E., Orioli, E., Pegoraro, A., Sangaletti, S., Portararo, P., Curti, A., Colombo, M.P., Di Virgilio, F., and Adinolfi, E. (2019). The P2X7 receptor modulates immune cells infiltration, ectonucleotidases expression and extracellular ATP levels in the tumor microenvironment. Oncogene 38, 3636– 3650. 10.1038/s41388-019-0684-y.

15. Chen, W., Wieraszko, A., Hogan, M. V., Yang, H.A., Kornecki, E., and Ehrlich, Y.H. (1996). Surface protein phosphorylation by ecto-protein kinase is required for the maintenance of hippocampal long-term potentiation. Proc. Natl. Acad. Sci. U. S. A. 93, 8688–8693. 10.1073/pnas.93.16.8688.

16. Yalak, G., Ehrlich, Y.H., and Olsen, B.R. (2014). Ecto-protein kinases and phosphatases: An emerging field for translational medicine. J. Transl. Med. 12, 1–6. 10.1186/1479-5876-12-165.

17. Bordoli, M.R., Yum, J., Breitkopf, S.B., Thon, J.N., Italiano, J.E., Xiao, J., Worby, C., Wong, S.K., Lin, G., Edenius, M., et al. (2014). A secreted tyrosine kinase acts in the extracellular environment. Cell 158, 1033–1044. 10.1016/j.cell.2014.06.048.

18. Rodríguez, F., Allende, C.C., and Allende, J.E. (2005). Protein kinase casein kinase 2 holoenzyme produced ectopically in human cells can be exported to the external side of the cellular membrane. Proc. Natl. Acad. Sci. U. S. A. 102, 4718– 4723. 10.1073/pnas.0501074102.

19. Jiang, H., Dong, J., Song, K., Wang, T., Huang, W., and Zhang, J. (2017). A novel allosteric site in casein kinase 2α discovered using combining bioinformatics and biochemistry methods. Nat. Publ. Gr. 38, 1691–1698. 10.1038/aps.2017.55.

20. Backe, S.J., Votra, S.B.D., Stokes, M.P., Sebestyén, E., Castelli, M., Torielli, L., Colombo, G., Woodford, M.R., Mollapour, M., and Bourboulia, D. (2023). PhosY-secretome profiling combined with kinase-substrate interaction screening defines active c-Src-driven extracellular signaling. Cell Rep. 42. 10.1016/j.celrep.2023.112539.

21. Sánchez-Pozo, J., Baker-Williams, A.J., Woodford, M.R., Bullard, R., Wei, B., Mollapour, M., Stetler-Stevenson, W.G., Bratslavsky, G., and Bourboulia, D. (2018). Extracellular Phosphorylation of TIMP-2 by Secreted c-Src Tyrosine Kinase Controls MMP-2 Activity. iScience 1, 87–96. 10.1016/j.isci.2018.02.004.

22. Tagliabracci, V.S., Wiley, S.E., Guo, X., Kinch, L.N., Durrant, E., Wen, J., Xiao, J., Cui, J., Nguyen, K.B., Engel, J.L., et al. (2015). A Single Kinase Generates the Majority of the Secreted Phosphoproteome. Cell 161, 1619–1632. 10.1016/j.cell.2015.05.028.

23. Tagliabracci, V.S., Pinna, L.A., and Dixon, J.E. (2013). Secreted protein kinases. Trends Biochem. Sci. 38, 121–130. 10.1016/j.tibs.2012.11.008.

24. Landesman-bollag, E., Romieu-mourez, È., Song, D.H., Sonenshein, G.E., Cardi, R.D., and Seldin, D.C. (2001). Protein kinase CK2 in mammary gland tumorigenesis. Oncogene 20, 3247–3257.

25. Trembley, J.H., Wang, G., Unger, G., Slaton, J., Ahmed, K., Medical, V.A., and Affairs, V. (2009). CK2: A key player in cancer biology. Cell Mol Life Sci 66, 1858– 1867. 10.1007/s00018-009-9154-y.CK2.

26. Bae, J.S., Park, S., Jamiyandorj, U., Kim, K.M., Noh, S.J., Kim, R., Sylvester, K.G., and Jang, K.Y. (2016). CK2a / CSNK2A1 Phosphorylates SIRT6 and Is Involved in the Progression of Breast Carcinoma and Predicts Shorter Survival of Diagnosed Patients. Am. J. Pathol. 186, 3297–3315. 10.1016/j.ajpath.2016.08.007.

27. Giusiano, S., Cochet, C., Filhol, O., Duchemin-pelletier, E., Bonnier, P., and Carcopino, X. (2010). Protein kinase CK2 a subunit over-expression correlates with metastatic risk in breast carcinomas_: Quantitative immunohistochemistry in tissue microarrays. Eur. J. Cancer 7, 0–9. 10.1016/j.ejca.2010.11.028.

28. Sajnaga, E., Kubiński, K., and Szyszka, R. (2013). Site-Directed Mutagenesis in the Research of Protein Kinases - The Case of Protein Kinase CK2. In Genetic Manipulation of DNA and Protein - Examples from Current Research.

29. Montenarh, M., and Götz, C. (2018). Ecto-Protein kinase CK2, the neglected form of CK2 (Review). Biomed. Reports 8, 307–313. 10.3892/br.2018.1069.

30. Issinger, O.G., Boldyreff, B., Brockel, C., and Pelton, J.J. (1992). Characterization of the α and β Subunits of Casein Kinase 2 by Far-UV CD Spectroscopy. Biochemistry 31, 6098–6103. 10.1021/bi00141a020.

31. Schnitzler, A., and Niefind, K. (2021). Structural basis for the design of bisubstrate inhibitors of protein kinase CK2 provided by complex structures with the substrate-competitive inhibitor heparin. Eur. J. Med. Chem. 214. 10.1016/j.ejmech.2021.113223.

32. Gatica, M., Jedlicki, A., Allende, C.C., and Allende, J.E. (1994). Activity of the E75E76 mutant of the α subunit of casein kinase II from Xenopus laevis. FEBS Lett. 339, 93–96. 10.1016/0014-5793(94)80392-7.

33. Hu, E., and Rubin, C.S. (1990). Expression of wild-type and mutated forms of the catalytic (α) subunit of Caenorhabditis elegans casein kinase II in Escherichia coli. J. Biol. Chem. 265, 20609–20615. 10.1016/s0021-9258(17)30546-x.

34. Weeks, A.M., Byrnes, J.R., Lui, I., and Wells, J.A. (2021). Mapping proteolytic neo-N termini at the surface of living cells. Proc. Natl. Acad. Sci. U. S. A. 118. 10.1073/pnas.2018809118.

35. Schaefer, K., Lui, I., Byrnes, J.R., Kang, E., Zhou, J., Weeks, A.M., and Wells, J.A. (2022). Direct Identification of Proteolytic Cleavages on Living Cells Using a Glycan-Tethered Peptide Ligase. ACS Cent. Sci. 8, 1447–1456. 10.1021/acscentsci.2c00899.

36. Pedram, K., Shon, D.J., Tender, G.S., Mantuano, N.R., Northey, J.J., Metcalf, K.J., Wisnovsky, S.P., Riley, N.M., Forcina, G.C., Malaker, S.A., et al. (2023). Design of a mucin-selective protease for targeted degradation of cancer-associated mucins. Nat. Biotechnol. 10.1038/s41587-023-01840-6.

37. Gray, M.A., Stanczak, M.A., Mantuano, N.R., Xiao, H., Pijnenborg, J.F.A., Malaker, S.A., Miller, C.L., Weidenbacher, P.A., Tanzo, J.T., Ahn, G., et al. (2020). Targeted glycan degradation potentiates the anticancer immune response in vivo. Nat. Chem. Biol. 16, 1376–1384. 10.1038/s41589-020-0622-x.

38. Hertz, N.T., Wang, B.T., Allen, J.J., Zhang, C., Dar, A.C., Burlingame, A.L., and Shokat, K.M. (2010). Chemical Genetic Approach for Kinase-Substrate Mapping by Covalent Capture of Thiophosphopeptides and Analysis by Mass Spectrometry. Curr. Protoc. Chem. Biol. 2, 15–36. 10.1002/9780470559277.ch090201.

39. Xu, H., Niu, M., Yuan, X., Wu, K., and Liu, A. (2020). CD44 as a tumor biomarker and therapeutic target. Exp. Hematol. Oncol., 1–14. 10.1186/s40164-020-00192-0.

40. Xu, H., Wu, K., Tian, Y., Liu, Q., and Han, N.A. (2016). CD44 correlates with clinicopathological characteristics and is upregulated by EGFR in breast cancer. Int. J. Oncol. 49, 1343–1350. 10.3892/ijo.2016.3639.

41. Zhang, H., Liang, X., Duan, C., Liu, C., and Zhao, Z. (2014). Galectin-3 as a Marker and Potential Therapeutic Target in Breast Cancer. PLoS One 9, 1–7. 10.1371/journal.pone.0103482.

42. Tao, J., Li, H., Li, Q., and Yang, Y. (2014). CD109 is a potential target for triple-negative breast cancer. Tumor Biol. 35, 12083–12090. 10.1007/s13277-014-2509-5.

43. Rodriguez, F.A., Contreras, C., and Allende, J.E. (2008). Protein kinase CK2 as an ectokinase_: The role of the regulatory CK2 _ subunit. Proc. Natl. Acad. Sci. 105, 5693–5698.

44. Siddiqui-Jain, A., Drygin, D., Streiner, N., Chua, P., Pierre, F., O’Brien, S.E., Bliesath, J., Omori, M., Huser, N., Ho, C., et al. (2010). CX-4945, an orally bioavailable selective inhibitor of protein kinase CK2, inhibits prosurvival and angiogenic signaling and exhibits antitumor efficacy. Cancer Res. 70, 10288– 10298. 10.1158/0008-5472.CAN-10-1893.

45. Kooreman, N.G., Kim, Y., Almeida, P.E. De, Levy, R., Davis, M.M., Wu, J.C., Kooreman, N.G., Kim, Y., Almeida, P.E. De, and Termglinchan, V. (2018). Autologous iPSC-Based Vaccines Elicit Anti-tumor Responses In Vivo Article Autologous iPSC-Based Vaccines Elicit Anti-tumor Responses In Vivo. Stem Cell 22, 501–513.e7. 10.1016/j.stem.2018.01.016.

46. Bishop, A.C., Buzko, O., and Shokat, K.M. (2001). Magic bullets for protein kinases. Trends Cell Biol. 11, 167–172. 10.1016/S0962-8924(01)01928-6.

47. Liu, Y., Shah, K., Yang, F., Witucki, L., and Shokat, K.M. (1998). Engineering Src family protein kinases with unnatural nucleotide specificity. Chem. Biol. 5, 91–101. 10.1016/S1074-5521(98)90143-0.

48. Agallou, M., and Karagouni, E. (2019). Detection of Antigen-specific T cells in Spleens of Vaccinated Mice Applying 3[H]-Thymidine Incorporation Assay and Luminex Multiple Cytokine Analysis Technology. Bio-protocol 9, 1–12. 10.21769/BioProtoc.3252.

49. Tender, G.S., and Bertozzi, C.R. (2023). Bringing enzymes to the proximity party. RSC Chem. Biol. 4, 986–1002. 10.1039/d3cb00084b.

50. Perez-Riverol, Y., Csordas, A., Bai, J., Bernal-Llinares, M., Hewapathirana, S., Kundu, D.J., Inuganti, A., Griss, J., Mayer, G., Eisenacher, M., et al. (2019). The PRIDE database and related tools and resources in 2019: Improving support for quantification data. Nucleic Acids Res. 47, D442–D450. 10.1093/nar/gky1106.

51. Hornsby, M., Paduch, M., Miersch, S., Sa, A., Matsuguchi, T., Lee, B., Wypisniak, K., Doak, A., King, D., Usatyuk, S., et al. (2015). A High Through-put Platform for Recombinant Antibodies to Folded Proteins * □. 2833–2847. 10.1074/mcp.O115.052209.

52. Zhang, J., Xin, L., Shan, B., Chen, W., Xie, M., Yuen, D., Zhang, W., Zhang, Z., Lajoie, G.A., and Ma, B. (2012). PEAKS DB_: De Novo Sequencing Assisted Database Search for Sensitive and Accurate Peptide Identification * □. 1–8. 10.1074/mcp.M111.010587.

53. Aizik, L., Dror, Y., Taussig, D., Barzel, A., Carmi, Y., and Wine, Y. (2021). Antibody Repertoire Analysis of Tumor-In fi ltrating B Cells Reveals Distinct Signatures and Distributions Across Tissues. 12, 1–13. 10.3389/fimmu.2021.705381.

54. Holton, O.D., Antibodies, M.-, Covell, D.G., Barbet, J., Sieber, S.M., Talley, M.J., and Weinstein, J.N. (2016). Biodistribution of monoclonal IgG1 , F ( ab ’) 2 , and Fab ’ in mice after intravenous injection. Comparison between anti-B cell ( anti-Lyb8 . 2 ) and irrelevant ( MOPC-21 ) antibodies.

55. Van Deventer, J.A., and Wittrup, K.D. (2014). Monoclonal Antibodies. In Monoclonal Antibodies: Methods and Protocols, pp. 151–181.

