## Supplemental Information for "Chemoproteomics reveals immunogenic and tumor-associated cell surface substrates of ectokinase CK2α"

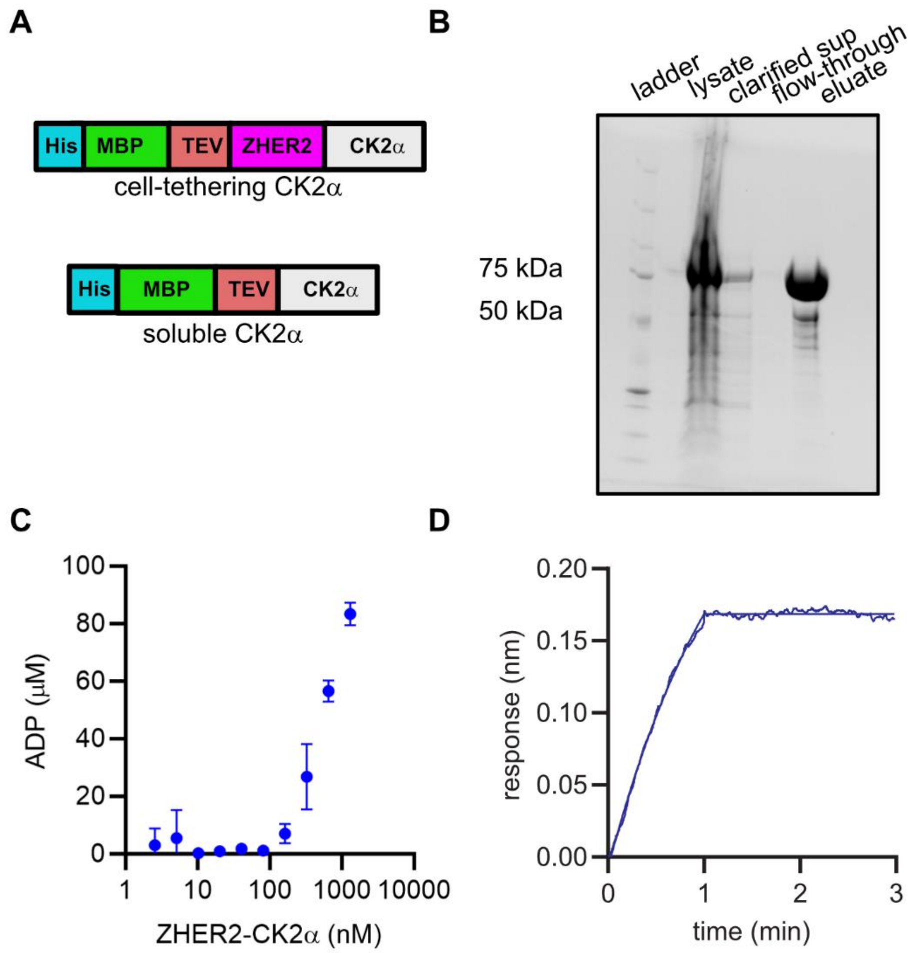


**Figure S1.** Expression of a ZHER2-CK2α fusion protein. (**A**) General diagram of CK2α constructs used. All constructs were expressed as fusions with N-terminal fusions to cleavable tags for purification. (**B**) ZHER2-CK2α is expressed cytosolically in E. coli as an N-terminal fusion to a maltose binding protein (MBP) tag, followed by a TEV-protease cut site. The MBP-fusions were stored in 10% glycerol at -80 ºC for long-term storage. The MBP was cleaved off by TEV protease treatment and removed according to manufacturer's protocols prior to subsequent experiments. (**C**) Kinase activity was measured against a commercial CK2α substrate peptide using Promega ADP-Glo universal kinase assay kit according to the manufacturer's instructions. (**D**) Binding of ZHER2-CK2α to recombinant HER2 was measured by biolayer interferometry. Association of soluble ZHER-CK2α to immobilized HER2-Fc-AviTag fusion begins at t = 0 min and dissociation begins at t = 1 min.

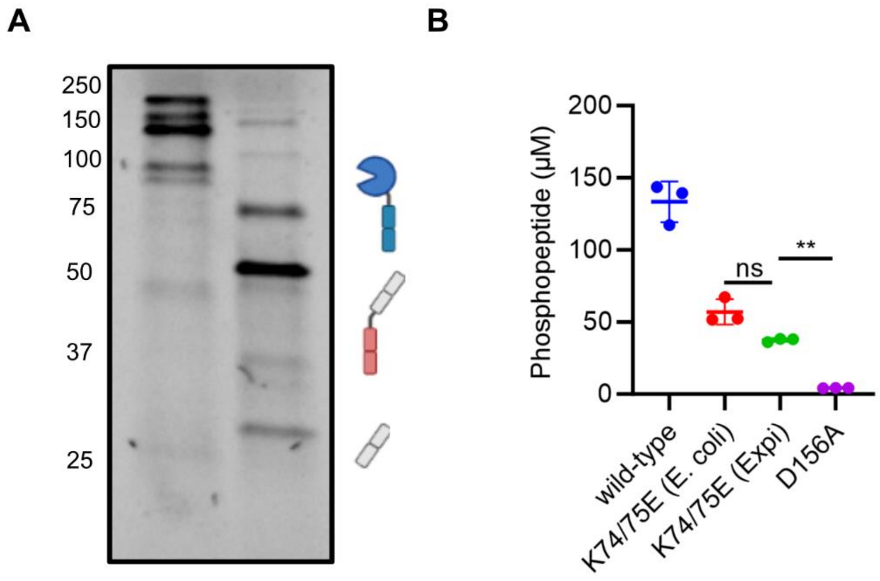


**Figure S2.** Expression and activity of CK2α as a knob-in-hole fusion with trastuzumab. (**A**) Co-transfection of a CK2α-Fc knob with trasutuzmab-hole and trastuzumab light chain in ExpiCHO cells affords an antibody-kinase fusion protein. CK2β is additionally transfected to improve secretion as it helps with expression but is not observed in purified products. (**B**) Kinase activity was measured against a commercial CK2α substrate peptide using Promega ADP-Glo universal kinase assay kit according to the manufacturer’s instructions.

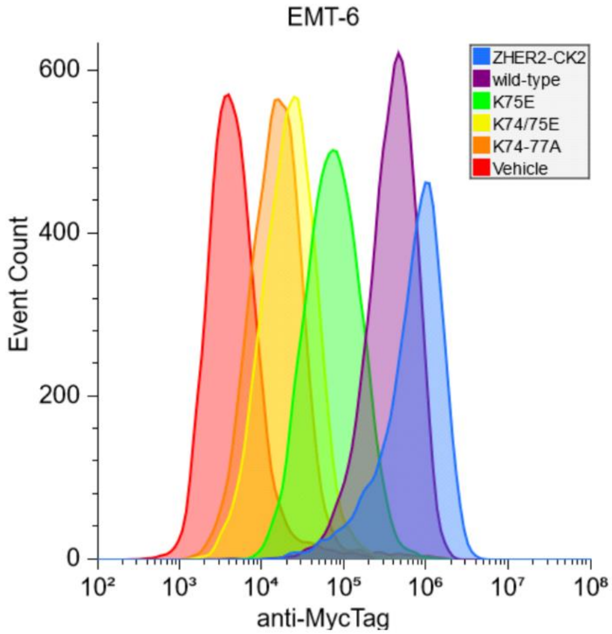


**Figure S3.** Representative flow histograms of CK2α-cMyc binding to hHER2-expressing EMT-6 murine breast cancer cells from **Figure 2B**. Purified recombinant ZHER2-CK2α or CK2α proteins, C-terminally fused to a cMyc peptide tag, were incubated with cells, and kinase binding was visualized with a fluorophore-conjugated anti-cMyc antibody by flow cytometry.

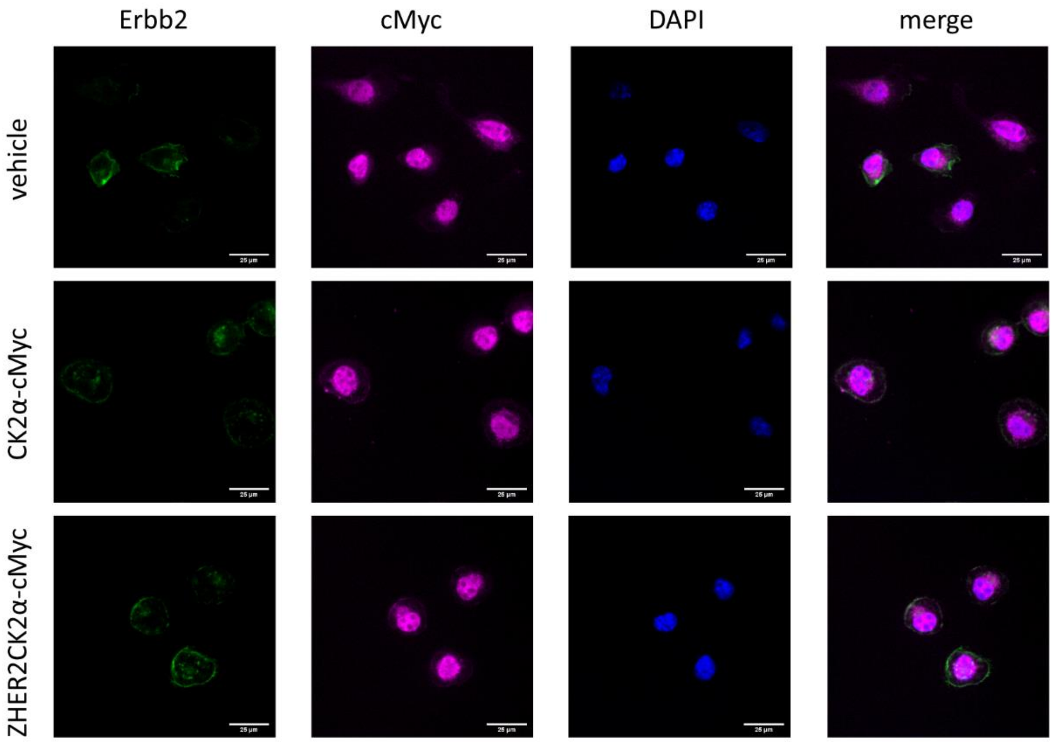


**Figure S4.** Microscopy images of CK2α binding to cell surfaces. EMT-6 cells were grown on glass slides in the presence of CK2α constructs C-terminally cMyc-tagged. Cells were fixed and permeabilized before being stained with DAPI, anti-Erbb2, and anti-cMyc (which also binds endogenous nuclear Myc). Pericellular localization is observed for Erbb2 and cMyc.

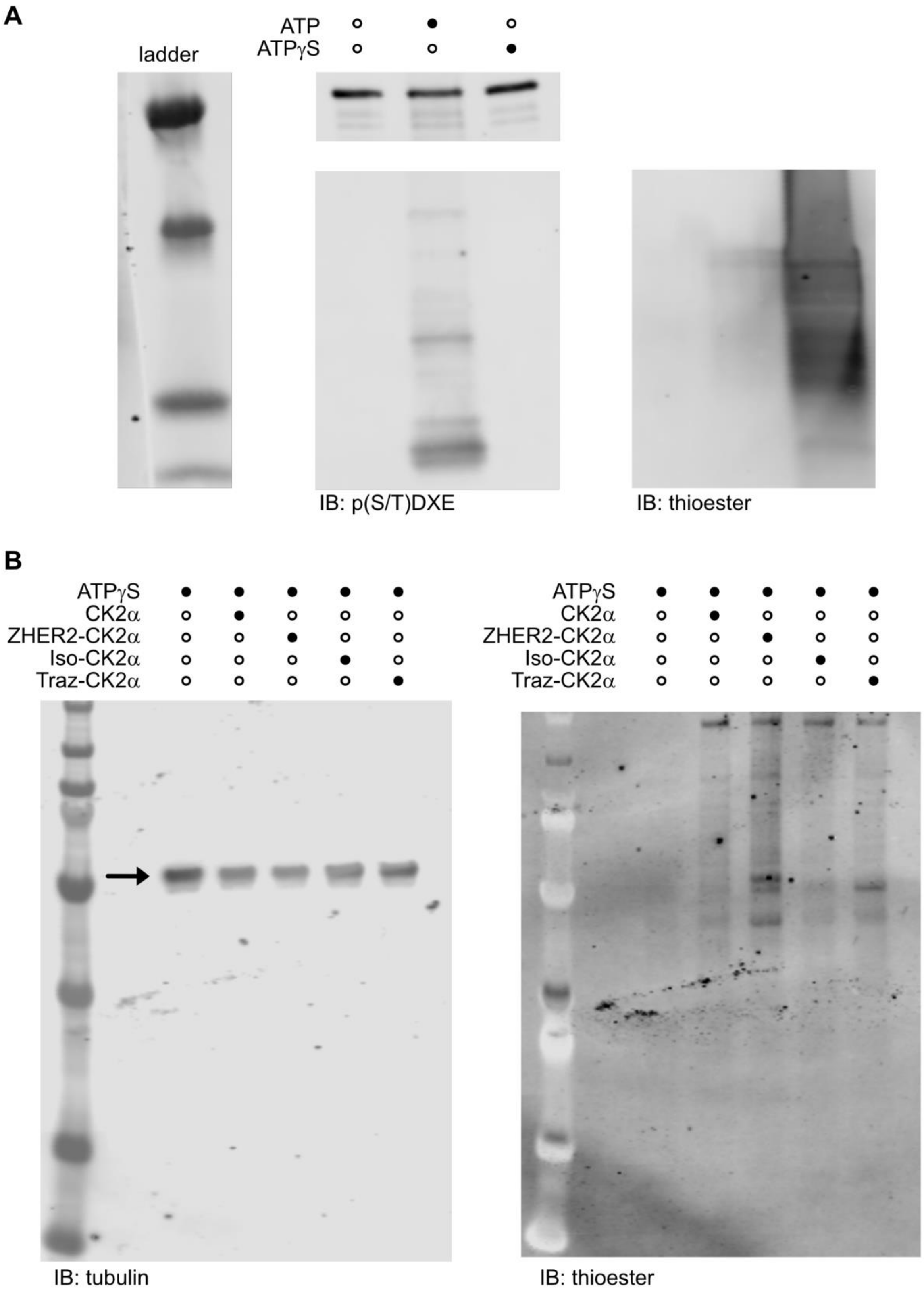
A
ladder
ATP
ATPγS
IB: p(S/T)DXE
IB: thioester
B
ATPγS
CK2α
ZHER2-CK2α
Iso-CK2α
Traz-CK2α
IB: tubulin
ATPγS
CK2α
ZHER2-CK2α
Iso-CK2α
Traz-CK2α
IB: thioester

**Figure S5.** Validation of CK2α activity with ATPγS and full Western blot from **Figure 2C**. (**A**) CK2α was incubated *in vitro* with purified substrate protein casein in the presence of ATP, ATPγS, or vehicle. Reactions were alkylated with p-nitrobenzylmesylate, separated by SDS-PAGE, transferred to PVDF membranes, and probed with either an anti-CK2 substrate antibody, anti-p(S/T)DXE, or anti-phosphothioester (thioester). (**B**) Full Western Blots of lysates from **Figure 2C**, with tubulin loading control (arrow), left, and thioester, right.

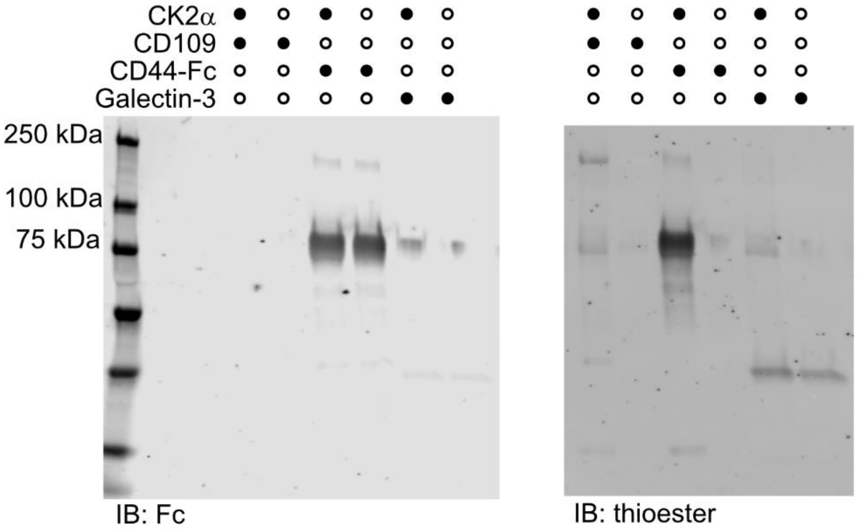


**Figure S6.** Full Western blot of **Figure 3C** with anti-Fc loading control (for CD44-Fc), left, and thioester, right.

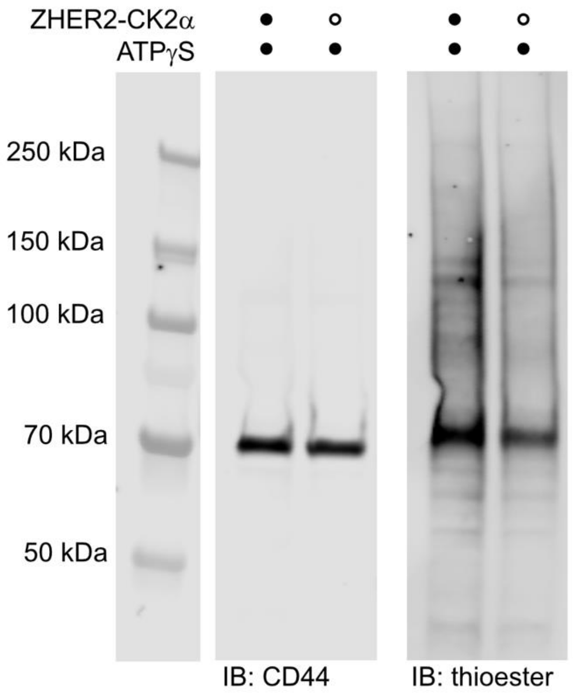


**Figure S7.** Full Western blot of **Figure 3D** with CD44, left, and thioester, right.

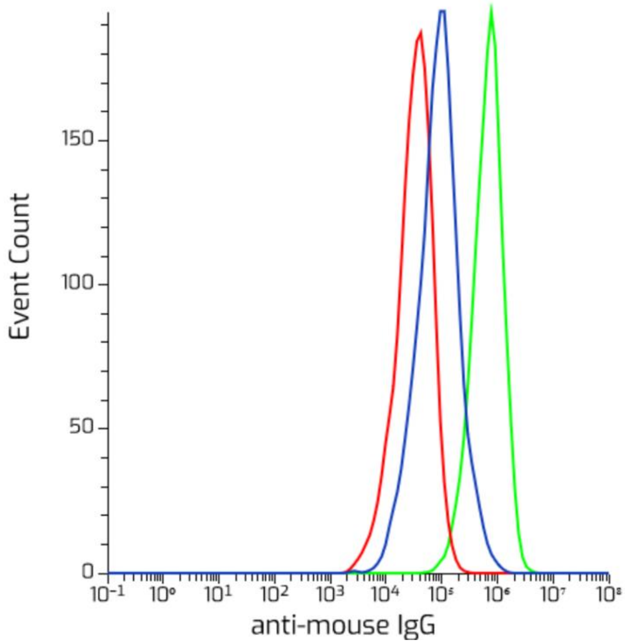


**Figure S8.** Representative flow cytograms for **Figure 4B**. hHER2-expressing EMT-6 cells were pretreated with cell-tethered CK2α (green) or vehicle (blue) before binding of Group 1 sera (1:50 dilution), followed by detection with an anti-mouse IgG AlexaFluor 647 secondary antibody by flow cytometry. Secondary antibody only on vehicle-treated cells was used as a control (red).

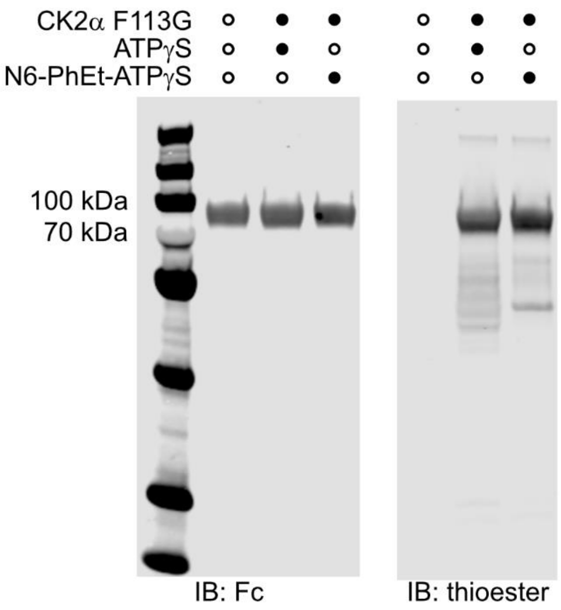


**Figure S9.** Gatekeeper mutation of CK2α F113G is permissive of N6-phenylethyl-ATP as a substrate. Recombinant murine CD44-Fc fusion protein was treated with or without CK2α F113G in the presence of either ATPγS or N6-phenylethyl-ATPγS (N6-PhEt-ATPγS). Reactions were alkylated with p-nitrobenzylmesylate, separated by SDS-PAGE, transferred to PVDF membranes, and probed with either anti-Fc, left, or anti-phosphothioester, right.

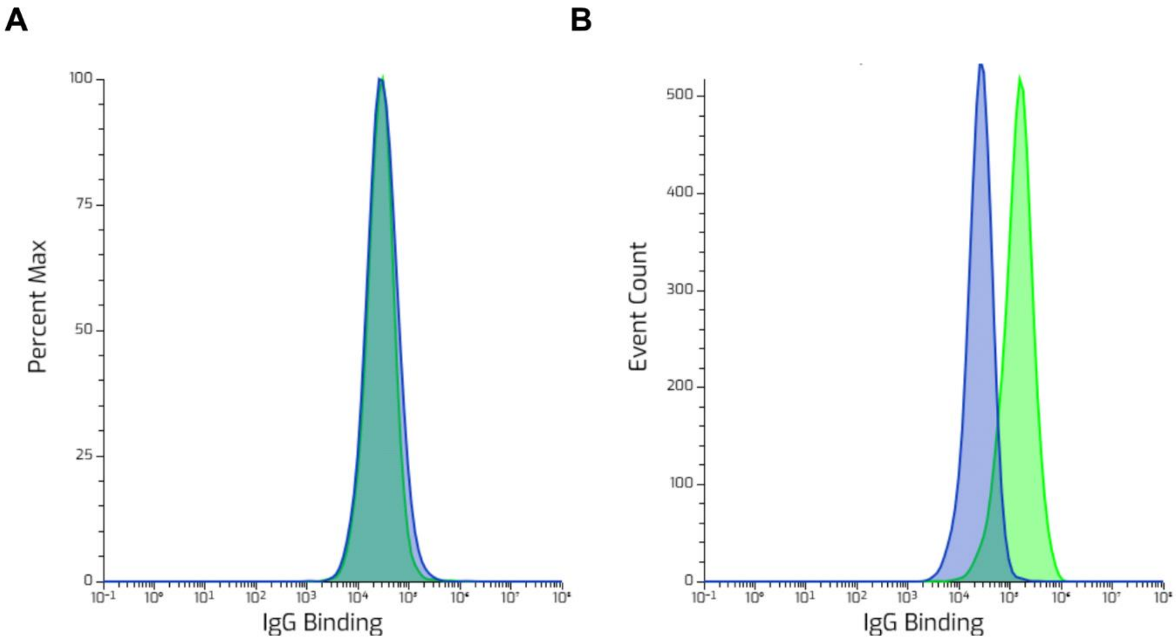


**Figure S10.** Representative flow cytograms of **Figure 4C**. (**A**,**B**) hHER2+ EMT-6 cells were pretreated with vehicle (**A**) or cell-tethered CK2α(F113G) (**B**) in the presence of 1 mM N6-phenylethyl-ATP before binding of Group 1 (green) or Group 2 (blue) sera (1:50 dilution), followed by detection with an anti-mouse IgG AlexaFluor 647 secondary antibody by flow cytometry.

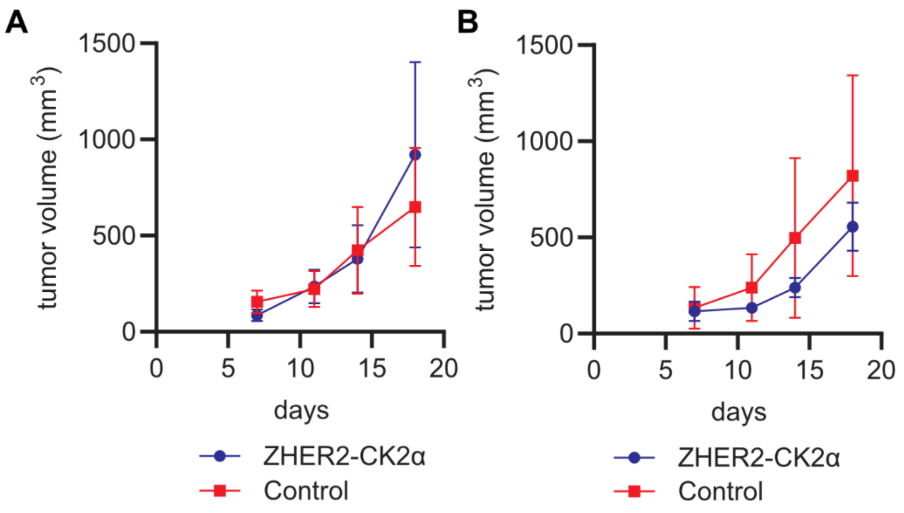


**Figure S11.** Group 2 and Group 3 tumor challenge experiments. (**A**) Mice were immunized weekly for four weeks with heat-killed hHER2-expressing EMT-6 cells. Mice were then challenged with live EMT-6 cells that were either hyperphosphorylated with cell-tethered CK2α or treated with vehicle *in vitro* prior to implantation. Averaged tumor growth curves from mice immunized with hyperphosphorylated EMT-6 cells. (**B**) Tumor challenge as in (**A**) but with naïve mice. For all groups, data show standard deviation of 5 mice per group.

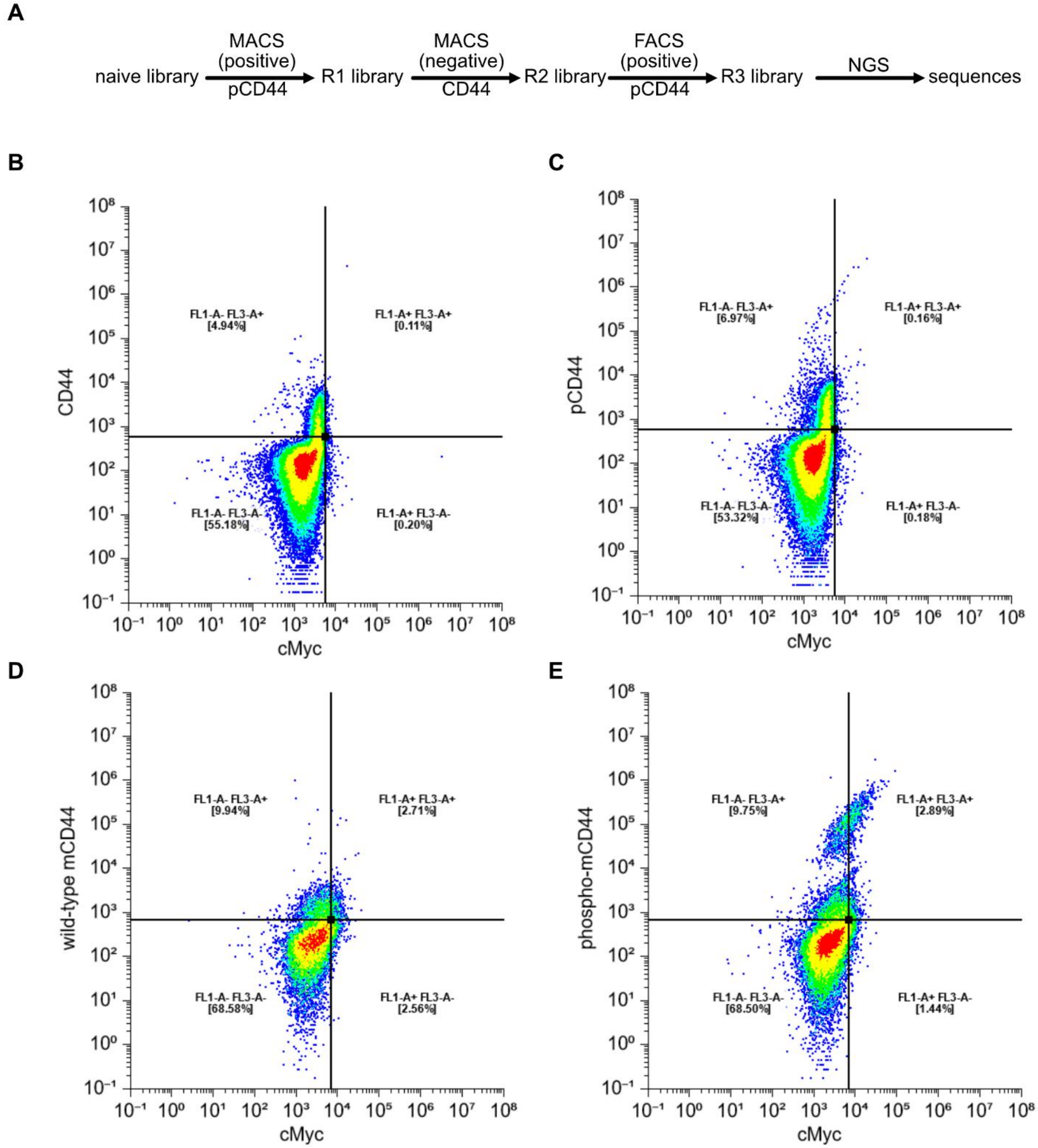


**Figure S12.** Yeast-display scheme and representative dot plots. (**A**) Splenocyte-derived VH sequences were subcloned into a common-light-chain yeast display vector and transformed into EBY100 yeast. Libraries were then subjected to successive rounds of magnetic sorting and fluorescence activated cell sorting. (**B,C**) Transformants from the parental library were tested for expression using an anti-cMyc tag antibody and antigen-specific binding to recombinant murine CD44[21-256]-Fc (**B**) or recombinant murine CD44[21-256]-Fc pretreated with CK2 *in vitro* (**C**). (**D, E**) The sorted library, subjected to three rounds of sorting with varying stringency, was tested for expression using an anti-cMyc tag antibody and antigen-specific binding to recombinant murine CD44[21-256]-Fc (**D**) or recombinant murine CD44[21-256]-Fc pretreated with CK2 *in vitro* (**E**).

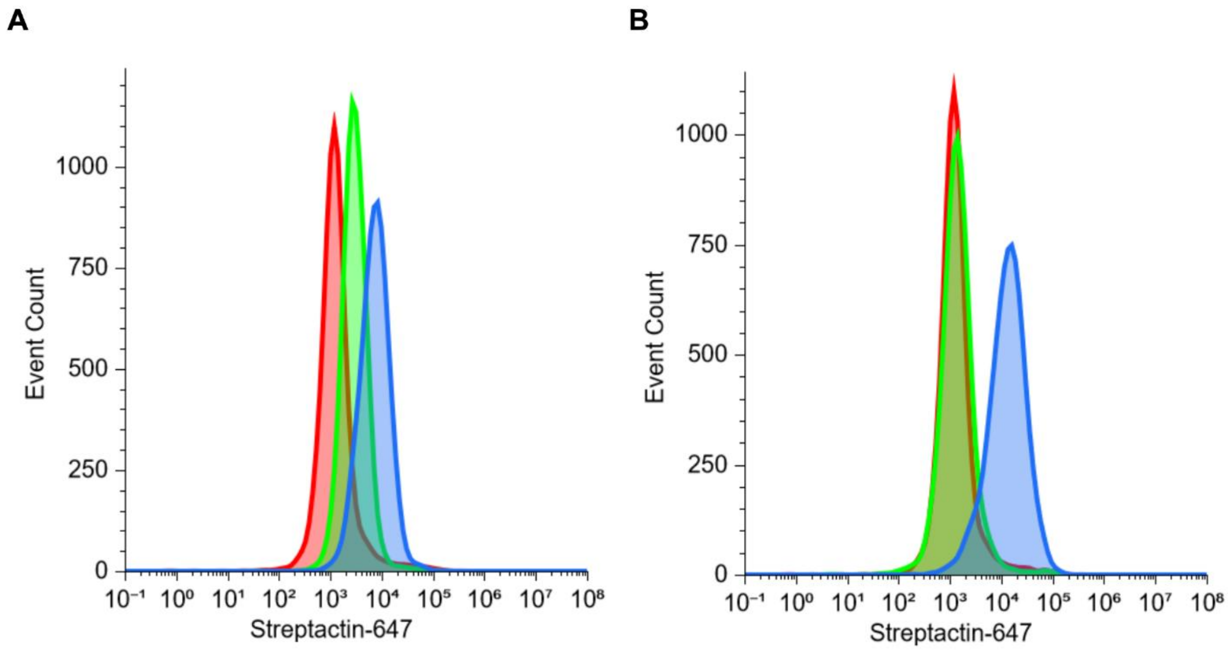


**Figure S13.** Representative histograms for **Figure 5D** for VH2 (**A**) and VH5 (**B**). Red represents Streptactin-647 alone. Green represents binding to untreated EMT-6 cells and Blue is binding to EMT-6 cells pretreated with cell-tethered CK2α.

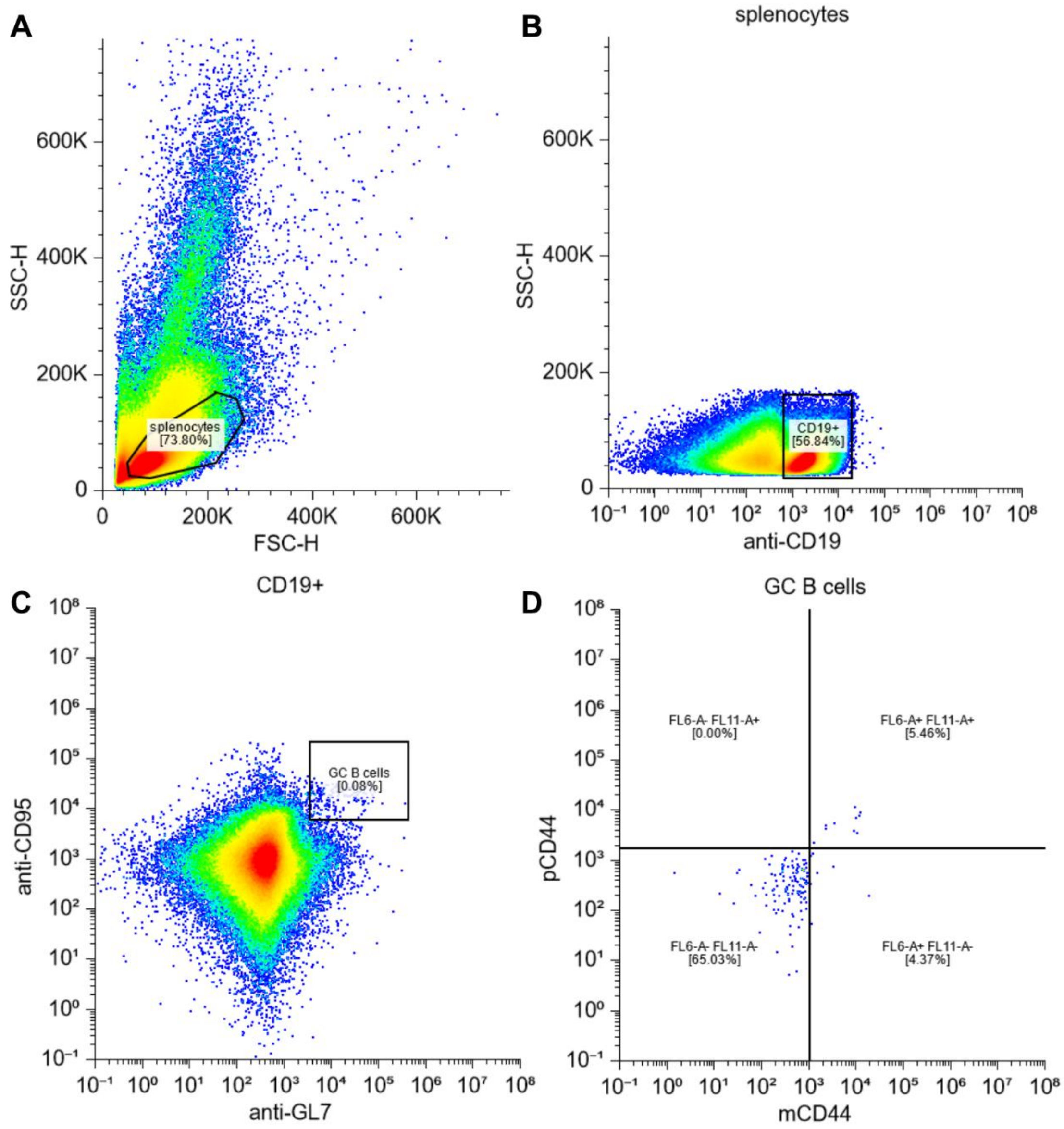


**Figure S14.** GC B cell gating scheme and representative dot plots for **Figure 5E**. Splenocytes were stained as described in the materials and methods and sorted for size (**A**), CD19 expression (**B**), and GC B cells were sorted based on into CD95 and GL7 expression (**C**). (**D**) Representative dot plot for GC B cell binding of mCD44 and pCD44.

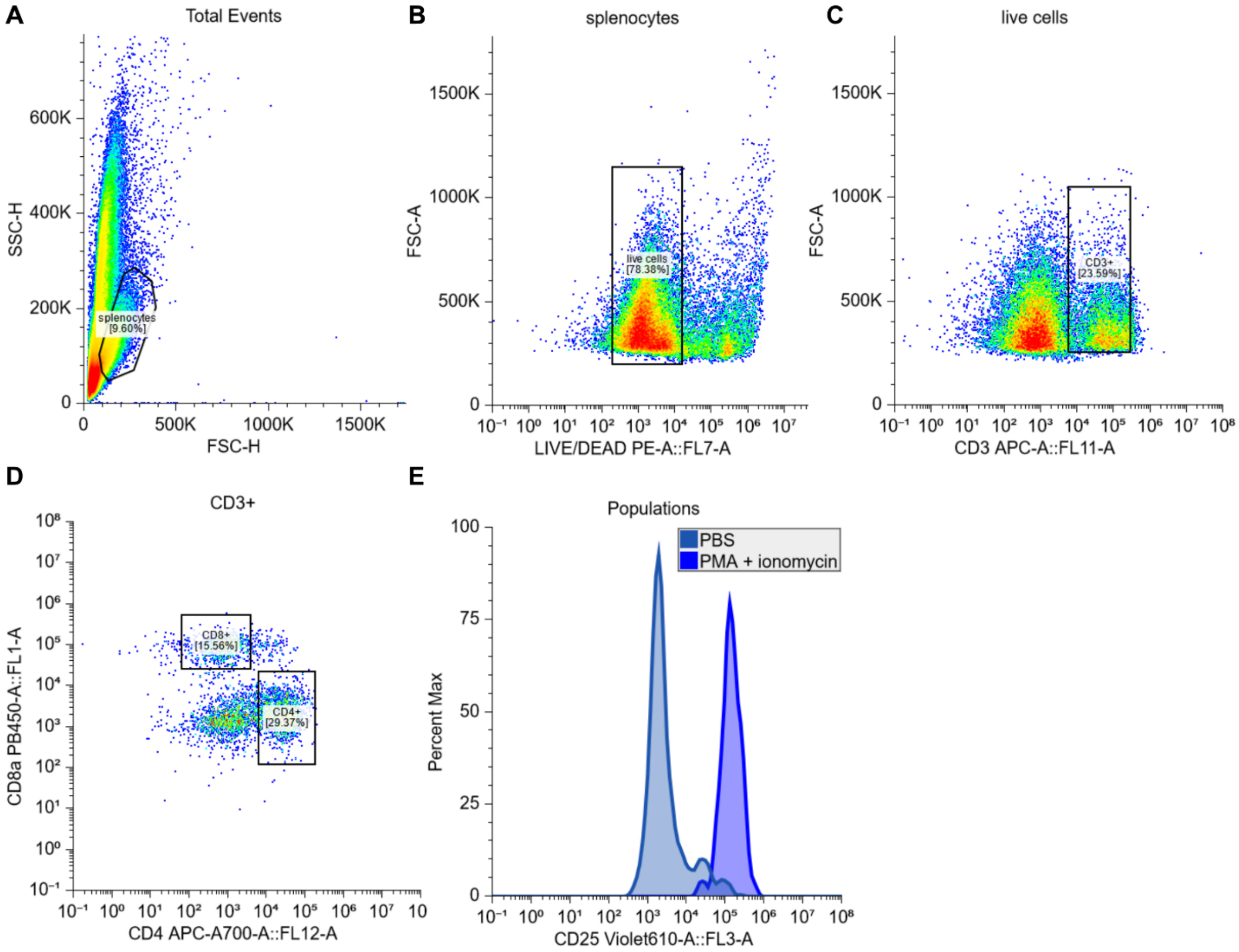


**Figure S15.** T cell gating schema and representative histograms for **Figure 5F-G**. Bulk splenocytes were treated as described in **Figure 5** and harvested with Versene. Splenocytes were stained as described in the materials and methods and sorted for size (**A**), viability (**B**), CD3 expression (**C**), and binned into CD4+ or CD8+ (**D**). (**E**) Representative data for CD25 expression on CD8+ T cells treated with either PBS or a mixture of PMA and ionomycin.

**Index of Supplementary Tables**

**Table S1.** Database search of EMT-6 derived ectophosphoproteome.

**Table S2.** LFQ results from EMT-6 derived ectophosphoproteome.

**Table S3.** Identification of CK2α phosphorylation sites on mCD44. Sample 16 corresponds to phosphorylated rmCD44. Sample 18 corresponds to rmCD44 treated with vehicle.

**Table S4.** Identification of CK2α phosphorylation sites on hCD44.

**Table S5.** Database search of SK-BR-3 derived ectophosphoproteome. Samples 1 and 2 were treated with CX-4945. Samples 3 and 4 were treated with DMSO.

**Table S6.** LFQ results from SK-BR-3 derived ectophosphoproteome. Samples 1 and 2 were treated with CX-4945. Samples 3 and 4 were treated with DMSO.

**Table S7.** Database search of immunoprecipitation of EMT-6 antigens. Animal IDs 886-890 correspond to Group 1 mice. Animal IDs 876-880 correspond to Group 2 mice.

**Table S8.** Oligonucleotide sequences

| name | sequence |
|---|---|
| CK2a K68A | AAGTTGTTGTTgccATTCTCAAGCCA |
| CK2a K68A/M REV | TTTCATTATTTGTGATGTTGATGGCTT |
| CK2a K68M | AAGTTGTTGTTAtgATTCTCAAGCCA |
| CK2a D156A | TGCACAGAGccGTCAAGCCC |
| CK2a D156A | TAATTCCCATGCTGTGACAATAATCC |
| CK2a I82C | AAAATTAAGCGTGAATGCAAGATTTTGGA |
| CK2a I82C | CTTCTTTTTTACTGGCTTGAGAATTTTAACAA |
| CK2a K75E | AAGCCAGTAAAAGAAAAGAAAATTAAGCGT |
| CK2a K75E | GAGAATTTTAACAACAACTTTTTCATTATTTGTGATG |
| CK2a FWD (pMAL C-term truncation cloning) | TAAggccgcgatatcgtc |
| CK2a[1-335] | ACCCATTCGAGCCTGGT |

| | |
|---|---|
| CK21[1-324] | GAAATAGGGGTGCTCCATTG |
| Ck2a-cMyc | tacctgcagggaatttattacaagtcttcttcggagataagcttttgttcagaacctccTGAACCgaggttctcccc |
| VH1 | CCC TCC TTT AAT TCC CGA KGT RMA GCT TCA GGA GTC |
| VH2 | CCC TCC TTT AAT TCC CGA GGT BCA GCT BCA GCA GTC |
| VH3 | CCC TCC TTT AAT TCC CCA GGT GCA GCT GAA GSA STC |
| VH4 | CCC TCC TTT AAT TCC CGA GGT CCA RCT GCA ACA RTC |
| VH5 | CCC TCC TTT AAT TCC CCA GGT YCA GCT BCA GCA RTC |
| VH6 | CCC TCC TTT AAT TCC CCA GGT YCA RCT GCA GCA GTC |
| VH7 | CCC TCC TTT AAT TCC CCA GGT CCA CGT GAA GCA GTC |
| VH8 | CCC TCC TTT AAT TCC CGA GGT GAA SST GGT GGA ATC |
| VH9 | CCC TCC TTT AAT TCC CGA VGT GAW GYT GGT GGA GTC |
| VH10 | CCC TCC TTT AAT TCC CGA GGT GCA GSK GGT GGA GTC |
| VH11 | CCC TCC TTT AAT TCC CGA KGT GCA MCT GGT GGA GTC |
| VH12 | CCC TCC TTT AAT TCC CGA GGT GAA GCT GAT GGA RTC |
| VH13 | CCC TCC TTT AAT TCC CGA GGT GCA RCT TGT TGA GTC |
| VH14 | CCC TCC TTT AAT TCC CGA RGT RAA GCT TCT CGA GTC |
| VH15 | CCC TCC TTT AAT TCC CGA AGT GAA RST TGA GGA GTC |
| VH16 | CCC TCC TTT AAT TCC CCA GGT TAC TCT RAA AGW GTS TG |
| VH17 | CCC TCC TTT AAT TCC CCA GGT CCA ACT VCA GCA RCC |
| VH18 | CCC TCC TTT AAT TCC CGA TGT GAA CTT GGA AGT GTC |
| VH19 | CCC TCC TTT AAT TCC CGA GGT GAA GGT CAT CGA GTC |
| H1 | GAG GAG AGA GAG AGA G CG AGG GGG AAG ACA TTT GGG |
| H2 | GAG GAG AGA GAG AGA G CC ARK GGA TAG ACH GAT GGG |
| VHrev | TCCCCCTCCGCCAGATCCTCCGCCTCCGCTggagacggtgaccagggttccttgacc |
| VH1fwd | gcttacccatacgatgttccagattacgctagRtYcagctgcaRcagtct |
| VH2fwd | gcttacccatacgatgttccagattacgctGAGtgcagctgaagSagtcagga |
| VH3fwd | gcttacccatacgatgttccagattacgctGAGgtgcagcttcaggagtcag |
| VH4fwd | gcttacccatacgatgttccagattacgctgaggtgaagcttctcgagtc |
| VH5fwd | gcttacccatacgatgttccagattacgctgaagtgaagctggtggagtc |
| VH6fwd | gcttacccatacgatgttccagattacgctggttactctgaaagagtc |
| VH7fwd | gcttacccatacgatgttccagattacgctgatccagttggtgcagt |
| VH8fwd | gcttacccatacgatgttccagattacgctgaagtgcagctgttggagac |
| VH9fwd | gcttacccatacgatgttccagattacgctcaggtgcagcttgtagagac |
| VH10fwd | gcttacccatacgatgttccagattacgctgaggttcagctgcagcagt |
| VH11fwd | gcttacccatacgatgttccagattacgctaggtgcagctggtgga |

Table S9. Plasmid DNA gene sequences and translations

| name | DNA sequence | translation |
|---|---|---|
| MBP-CK2α | atgaaaatccaccatcaccaccaccacgaagaaggtaaactggtaatctggattaa<br>cggcgataaaggctataacggtctcgctgaagtcggtaagaaattcgagaaagata<br>ccggaattaaagtcaccgttgagcatccggataaactggaagagaaattcccacag<br>gttgcggcaactggcgatggccctgacattatcttctgggcacacgaccgctttgg<br>tggctacgctcaatctggcctgttggctgaaatcaccccggacaaagcgttccagg<br>acaagctgtatccgtttacctgggatgccgtacgttacaacggcaagctgattgct<br>tacccgatcgctgttgaagcgttatcgctgatttataacaaagatctgctgccgaa | MKIHHHHHHEEGKLVIWINGDKGYNGL<br>AEVGKKFEKDTGIKVTVEHPDKLEEKF<br>PQVAATGDGPDIIFWAHDRFGGYAQSG<br>LLAEITPDKAFQDKLYPFTWDAVRYNG<br>KLIAYPIAVEALSLIYNKDLLPNPPKT<br>WEEIPALDKELKAKGKSALMFNLQEPY<br>FTWPLIAADGGYAFKYENGKYDIKDVG |

| | | |
|---|---|---|
| | cccgccaaaaacctgggaagagatcccggcgctggataaagaactgaaagcgaaag<br>gtaagagcgcgctgatgttcaacctgcaagaaccgtacttcacctggccgctgatt<br>gctgctgacgggggttatgcgttcaagtatgaaaacggcaagtacgacattaaaga<br>cgtgggcgtggataacgctggcgcgaaagcgggtctgaccttcctggttgacctga<br>ttaaaaacaaacacatgaatgcagacaccgattactccatcgcagaagcagccttt<br>aataaaggcgaaacagcgatgaccatcaacggcccgtgggcatggtccaacatcga<br>caccagcaaagtgaattatggtgtaacggtactgccgaccttcaagggtcaaccat<br>ccaaaccgttcgttggcgtgctgagcgcaggtattaacgccgccagtccgaacaaa<br>gagctggcaaaagagttcctcgaaaactatctgctgactgatgaaggtctggaagc<br>ggttaataaagacaaaccgctgggtgccgtagcgctgaagtcttacgaggaagagt<br>tggtgaaagatccgcgtattgccgccactatggaaaacgcccagaaaggtgaaatc<br>atgccgaacatcccgcagatgtccgctttctggtatgccgtgcgtactgcggtgat<br>caacgccgccagcggtcgtcagactgtcgatgaagccctgaaagacgcgcagacta<br>attcgagctcgaacaacaacaacaataacaataacaacaacctcggggagaacctg<br>tacttccagatgctgatgggcggccgcATGTCGGGACCCGTGCCAAGCAGGGCCAG<br>AGTTTACACAGATGTTAATACACACAGACCTCGAGAATACTGGGATTACGAGTCAC<br>ATGTGGTGGAATGGGGAAATCAAGATGACTACCAGCTGGTTCGAAAATTAGGCCGA<br>GGTAAATACAGTGAAGTATTTGAAGCCATCAACATCACAAATAATGAAAAAGTTGT<br>TGTTAAAATTCTCAAGCCAGTAAAAAAGAAGAAAATTAAGCGTGAAATAAAGATTT<br>TGGAGAATTTGAGAGGAGGTCCCAACATCATCACACTGGCAGACATTGTAAAAGAC<br>CCTGTGTCACGAACCCCCGCCTTGGTTTTTGAACACGTAAACAACACAGACTTCAA<br>GCAATTGTACCAGACGTTAACAGACTATGATATTCGATTTTACATGTATGAGATTC<br>TGAAGGCCCTGGATTATTGTCACAGCATGGGAATTATGCACAGAGATGTCAAGCCC<br>CATAATGTCATGATTGATCATGAGCACAGAAAGCTACGACTAATAGACTGGGGTTT<br>GGCTGAGTTTTATCATCCTGGCCAAGAATATAATGTCCGAGTTGCTTCCCGATACT<br>TCAAAGGTCCTGAGCTACTTGTAGACTATCAGATGTACGATTATAGTTTGGATATG<br>TGGAGTTTGGGTTGTATGCTGGCAAGTATGATCTTTCGGAAGGAGCCATTTTTCCA<br>TGGACATGACAATTATGATCAGTTGGTGAGGATAGCCAAGGTTCTGGGGACAGAAG<br>ATTTATATGACTATATTGACAAATACAACATTGAATTAGATCCACGTTTCAATGAT<br>ATCTTGGGCAGACACTCTCGAAAGCGATGGGAACGCTTTGTCCACAGTGAAAATCA<br>GCACCTTGTCAGCCCTGAGGCCTTGGATTTCCTGGACAAACTGCTGCGATATGACC<br>ACCAGTCACGGCTTACTGCAAGAGAGGCAATGGAGCACCCCTATTTCTACACTGTT<br>GTGAAGGACCAGGCTCGAATGGGTTCATCTAGCATGCCAGGGGGCAGTACGCCCGT<br>CAGCAGCGCCAATATGATGTCAGGGATTTCTTCAGTGCCAACCCCTTCACCCCTTG<br>GACCTCTGGCAGGCTCACCAGTGATTGCTGCTGCCAACCCCCTTGGGATGCCTGTT<br>CCAGCTGCCGCTGGCGCTCAGCAG | VDNAGAKAGLTFLVDLIKNKHMNADTD<br>YSIAEAAFNKGETAMTINGPWAWSNID<br>TSKVNYGVTVLPTFKGQPSKPFVGVLS<br>AGINAASPNKELAKEFLENYLLTDEGL<br>EAVNKDKPLGAVALKSYEEELVKDPRI<br>AATMENAQKGEIMPNIPQMSAFWYAVR<br>TAVINAASGRQTVDEALKDAQTNSSSN<br>NNNNNNNNNLGENLYFQMLMGGRMSGP<br>VPSRARVYTDVNTHRPREYWDYESHVV<br>EWGNQDDYQLVRKLGRGKYSEVFEAIN<br>ITNNEKVVVKILKPVKKKKIKREIKIL<br>ENLRGGPNIITLADIVKDPVSRTPALV<br>FEHVNNTDFKQLYQTLTDYDIRFYMYE<br>ILKALDYCHSMGIMHRDVKPHNVMIDH<br>EHRKLRLIDWGLAEFYHPGQEYNVRVA<br>SRYFKGPELLVDYQMYDYSLDMWSLGC<br>MLASMIFRKEPFFHGHDNYDQLVRIAK<br>VLGTEDLYDYIDKYNIELDPRFNDILG<br>RHSRKRWERFVHSENQHLVSPEALDFL<br>DKLLRYDHQSRLTAREAMEHPYFYTVV<br>KDQARMGSSSMPGGSTPVSSANMMSGI<br>SSVPTPSPLGPLAGSPVIAAANPLGMP<br>VPAAAGAQQ |
| MBP-ZHER2-CK2α [1-336] | atgaaaatccaccatcaccaccaccacgaagaaggtaaactggtaatctggattaa<br>cggcgataaaggctataacggtctcgctgaagtcggtaagaaattcgagaaagata<br>ccggaattaaagtcaccgttgagcatccggataaactggaagagaaattcccacag<br>gttgcggcaactggcgatggccctgacattatcttctgggcacacgaccgctttgg<br>tggctacgctcaatctggcctgttggctgaaatcaccccggacaaagcgttccagg<br>acaagctgtatccgtttacctgggatgccgtacgttacaacggcaagctgattgct<br>tacccgatcgctgttgaagcgttatcgctgatttataacaaagatctgctgccgaa<br>cccgccaaaaacctgggaagagatcccggcgctggataaagaactgaaagcgaaag<br>gtaagagcgcgctgatgttcaacctgcaagaaccgtacttcacctggccgctgatt<br>gctgctgacgggggttatgcgttcaagtatgaaaacggcaagtacgacattaaaga<br>cgtgggcgtggataacgctggcgcgaaagcgggtctgaccttcctggttgacctga<br>ttaaaaacaaacacatgaatgcagacaccgattactccatcgcagaagcagccttt<br>aataaaggcgaaacagcgatgaccatcaacggcccgtgggcatggtccaacatcga<br>caccagcaaagtgaattatggtgtaacggtactgccgaccttcaagggtcaaccat<br>ccaaaccgttcgttggcgtgctgagcgcaggtattaacgccgccagtccgaacaaa<br>gagctggcaaaagagttcctcgaaaactatctgctgactgatgaaggtctggaagc<br>ggttaataaagacaaaccgctgggtgccgtagcgctgaagtcttacgaggaagagt<br>tggtgaaagatccgcgtattgccgccactatggaaaacgcccagaaaggtgaaatc<br>atgccgaacatcccgcagatgtccgctttctggtatgccgtgcgtactgcggtgat<br>caacgccgccagcggtcgtcagactgtcgatgaagccctgaaagacgcgcagacta<br>attcgagctcgaacaacaacaacaataacaataacaacaacctcggggagaacctg<br>tacttccagatgctgatgggcGTGGATAACAAATTTAACAAAGAAATGCGCAACGC<br>GTATTGGGAAATTGCGCTGCTGCCGAACCTGAACAACCAGCAGAAACGCGCGTTTA<br>TTCGCAGCCTGTATGATGATCCGAGCCAGAGCGCGAACCTGCTGGCGGAAGCGAAA<br>AAACTGAACGATGCGCAGGCGCCGAAAggcggtggcggaagtggcggcggtggaac<br>tagtggccgcATGTCGGGACCCGTGCCAAGCAGGGCCAGAGTTTACACAGATGTTA<br>ATACACACAGACCTCGAGAATACTGGGATTACGAGTCACATGTGGTGGAATGGGGA<br>AATCAAGATGACTACCAGCTGGTTCGAAAATTAGGCCGAGGTAAATACAGTGAAGT<br>ATTTGAAGCCATCAACATCACAAATAATGAAAAAGTTGTTGTTAAAATTCTCAAGC<br>CAGTAAAAAAGAAGAAAATTAAGCGTGAAATAAAGATTTTGGAGAATTTGAGAGGA<br>GGTCCCAACATCATCACACTGGCAGACATTGTAAAAGACCCTGTGTCACGAACCCC<br>CGCCTTGGTTTTTGAACACGTAAACAACACAGACTTCAAGCAATTGTACCAGACGT<br>TAACAGACTATGATATTCGATTTTACATGTATGAGATTCTGAAGGCCCTGGATTAT<br>TGTCACAGCATGGGAATTATGCACAGAGATGTCAAGCCCCATAATGTCATGATTGA<br>TCATGAGCACAGAAAGCTACGACTAATAGACTGGGGTTTGGCTGAGTTTTATCATC | MKIHHHHHHEEGKLVIWINGDKGYNGL<br>AEVGKKFEKDTGIKVTVEHPDKLEEKF<br>PQVAATGDGPDIIFWAHDRFGGYAQSG<br>LLAEITPDKAFQDKLYPFTWDAVRYNG<br>KLIAYPIAVEALSLIYNKDLLPNPPKT<br>WEEIPALDKELKAKGKSALMFNLQEPY<br>FTWPLIAADGGYAFKYENGKYDIKDVG<br>VDNAGAKAGLTFLVDLIKNKHMNADTD<br>YSIAEAAFNKGETAMTINGPWAWSNID<br>TSKVNYGVTVLPTFKGQPSKPFVGVLS<br>AGINAASPNKELAKEFLENYLLTDEGL<br>EAVNKDKPLGAVALKSYEEELVKDPRI<br>AATMENAQKGEIMPNIPQMSAFWYAVR<br>TAVINAASGRQTVDEALKDAQTNSSSN<br>NNNNNNNNNLGENLYFQMLMGVDNKFN<br>KEMRNAYWEIALLPNLNNQQKRAFIRS<br>LYDDPSQSANLLAEAKKLNDAQAPKGG<br>GGSGGGGTSGRMSGPVPSRARVYTDVN<br>THRPREYWDYESHVVEWGNQDDYQLVR<br>KLGRGKYSEVFEAINITNNEKVVVKIL<br>KPVKKKKIKREIKILENLRGGPNIITL<br>ADIVKDPVSRTPALVFEHVNNTDFKQL<br>YQTLTDYDIRFYMYEILKALDYCHSMG<br>IMHRDVKPHNVMIDHEHRKLRLIDWGL<br>AEFYHPGQEYNVRVASRYFKGPELLVD<br>YQMYDYSLDMWSLGCMLASMIFRKEPF<br>FHGHDNYDQLVRIAKVLGTEDLYDYID<br>KYNIELDPRFNDILGRHSRKRWERFVH<br>SENQHLVSPEALDFLDKLLRYDHQSRL<br>TAREAMEHPYFYTVVKDQARMGS |

| | | |
|---|---|---|
| | CTGGCCAAGAATATAATGTCCGAGTTGCTTCCCGATACTTCAAAGGTCCTGAGCTA<br>CTTGTAGACTATCAGATGTACGATTATAGTTTGGATATGTGGAGTTTGGGTTGTAT<br>GCTGGCAAGTATGATCTTTCGGAAGGAGCCATTTTTCCATGGACATGACAATTATG<br>ATCAGTTGGTGAGGATAGCCAAGGTTCTGGGGACAGAAGATTTATATGACTATATT<br>GACAAATACAACATTGAATTAGATCCACGTTTCAATGATATCTTGGGCAGACACTC<br>TCGAAAGCGATGGGAACGCTTTGTCCACAGTGAAAATCAGCACCTTGTCAGCCCTG<br>AGGCCTTGGATTTCCTGGACAAACTGCTGCGATATGACCACCAGTCACGGCTTACT<br>GCAAGAGAGGCAATGGAGCACCCCTATTTCTACACTGTTGTGAAGGACCAGGCTCG<br>AATGGGTTCA | |
| CK2α[1-336]-Fc knob | ATGTACAGGATGCAACTCCTGTCTTGCATTGCACTAAGTCTTGCACTTGTCACGAA<br>TTCGATGTCGGGACCCGTGCCAAGCAGGGCCAGAGTTTACACAGATGTTAATACAC<br>ACAGACCTCGAGAATACTGGGATTACGAGTCACATGTGGTGGAATGGGGAAATCAA<br>GATGACTACCAGCTGGTTCGAAAATTAGGCCGAGGTAAATACAGTGAAGTATTTGA<br>AGCCATCAACATCACAAATAATGAAAAAGTTGTTGTTAAAATTCTCAAGCCAGTAA<br>AAAAGAAGAAAATTAAGCGTGAAATAAAGATTTTGGAGAATTTGAGAGGAGGTCCC<br>AACATCATCACACTGGCAGACATTGTAAAAGACCCTGTGTCACGAACCCCCGCCTT<br>GGTTTTTGAACACGTAAACAACACAGACTTCAAGCAATTGTACCAGACGTTAACAG<br>ACTATGATATTCGATTTTACATGTATGAGATTCTGAAGGCCCTGGATTATTGTCAC<br>AGCATGGGAATTATGCACAGAGATGTCAAGCCCCATAATGTCATGATTGATCATGA<br>GCACAGAAAGCTACGACTAATAGACTGGGGTTTGGCTGAGTTTTATCATCCTGGCC<br>AAGAATATAATGTCCGAGTTGCTTCCCGATACTTCAAAGGTCCTGAGCTACTTGTA<br>GACTATCAGATGTACGATTATAGTTTGGATATGTGGAGTTTGGGTTGTATGCTGGC<br>AAGTATGATCTTTCGGAAGGAGCCATTTTTCCATGGACATGACAATTATGATCAGT<br>TGGTGAGGATAGCCAAGGTTCTGGGGACAGAAGATTTATATGACTATATTGACAAA<br>TACAACATTGAATTAGATCCACGTTTCAATGATATCTTGGGCAGACACTCTCGAAA<br>GCGATGGGAACGCTTTGTCCACAGTGAAAATCAGCACCTTGTCAGCCCTGAGGCCT<br>TGGATTTCCTGGACAAACTGCTGCGATATGACCACCAGTCACGGCTTACTGCAAGA<br>GAGGCAATGGAGCACCCCTATTTCTACACTGTTGTGAAGGACCAGGCTCGAATGGG<br>TACCGGTTCTGGTGGTGGTGGTCTTGAGGTTCTCTTTCAGGGGCCGAGCTCGGGCG<br>GAGGATCGGGTGGAGGCgagcccaaatcttgtgacaaaactcacacatgcCCCCCC<br>TGCCCAGCGCCAGAATTGCTGGGCGGACCCAGCGTGTTCCTGTTCCCCCCCAAACC<br>TAAAGACACCCTGATGATCAGCCGAACCCCTGAGGTGACCTGCGTGGTGGTGGACG<br>TGAGCCACGAGGACCCCGAGGTGAAGTTCAACTGGTATGTGGACGGCGTGGAGGTC<br>CACAATGCCAAAACGAAGCCCAGGGAGGAGCAGTACAACAGCACCTACAGGGTAGT<br>GAGCGTCTTGACCGTGCTGCACCAGGACTGGCTGAACGGCAAGGAATACAAATGCA<br>AGGTCAGCAATAAGGCTCTGCCGGCTCCTATCGAGAAGACAATCAGCAAGGCAAAG<br>GGCCAGCCACGCGAACCGCAGGTGTATACTCTGCCCCCCAGCCGGGACGAGCTGAC<br>CAAGAACCAGGTGTCCCTGTGGTGTCTGGTGAAAGGCTTCTACCCCAGCGACATCG<br>CTGTGGAGTGGGAGAGTAACGGGCAGCCCGAGAACAACTACAAGACCACGCCTCCT<br>GTGCTGGACAGCGACGGCAGCTTCTTCCTGTATAGCAAGCTCACCGTGGACAAGAG<br>CAGGTGGCAACAGGGCAACGTGTTCAGCTGCTCTGTGATGCACGAGGCCCTGCACA<br>ACCATTACACCCAGAAGAGTCTCAGTCTGAGCCCGGGAAAGGGTGGCTCTCATCAT<br>CACCATCACCAC | MYRMQLLSCIALSLALVTNSMSGPVPS<br>RARVYTDVNTHRPREYWDYESHVVEWG<br>NQDDYQLVRKLGRGKYSEVFEAINITN<br>NEKVVVKILKPVKKKKIKREIKILENL<br>RGGPNIITLADIVKDPVSRTPALVFEH<br>VNNTDFKQLYQTLTDYDIRFYMYEILK<br>ALDYCHSMGIMHRDVKPHNVMIDHEHR<br>KLRLIDWGLAEFYHPGQEYNVRVASRY<br>FKGPELLVDYQMYDYSLDMWSLGCMLA<br>SMIFRKEPFFHGHDNYDQLVRIAKVLG<br>TEDLYDYIDKYNIELDPRFNDILGRHS<br>RKRWERFVHSENQHLVSPEALDFLDKL<br>LRYDHQSRLTAREAMEHPYFYTVVKDQ<br>ARMGTGSGGGGLEVLFQGPSSGGGSGG<br>GEPKSCDKTHTCPPCPAPELLGGPSVF<br>LFPPKPKDTLMISRTPEVTCVVVDVSH<br>EDPEVKFNWYVDGVEVHNAKTKPREEQ<br>YNSTYRVVSVLTVLHQDWLNGKEYKCK<br>VSNKALPAPIEKTISKAKGQPREPQVY<br>TLPPSRDELTKNQVSLWCLVKGFYPSD<br>IAVEWESNGQPENNYKTTPPVLDSDGS<br>FFLYSKLTVDKSRWQQGNVFSCSVMHE<br>ALHNHYTQKSLSLSPGKGGSHHHHHH |
| CK2β | ATGCGAATGCAGCTGCTGCTGCTGATTGCGCTGAGCCTGGCGCTGGTGACCAACAG<br>CGCTAGCCACCACCATCATCACCACCATCACGGATCCAGCAGCTCAGAGGAGGTGT<br>CCTGGATTTCCTGGTTCTGTGGGCTCCGTGGCAATGAATTCTTCTGTGAAGTGGAT<br>GAAGACTACATCCAGGACAAATTTAATCTTACTGGACTCAATGAGCAGGTCCCTCA<br>CTATCGACAAGCTCTAGACATGATCTTGGACCTGGAGCCTGATGAAGAACTGGAAG<br>ACAACCCCAACCAGAGTGACCTGATTGAGCAGGCAGCCGAGATGCTTTATGGATTG<br>ATCCACGCCCGCTACATCCTTACCAACCGTGGCATCGCCCAGATGTTGGAAAAGTA<br>CCAGCAAGGAGACTTTGGTTACTGTCCTCGTGTGTACTGTGAGAACCAGCCAATGC<br>TTCCCATTGGCCTTTCAGACATCCCAGGTGAAGCCATGGTGAAGCTCTACTGCCCC<br>AAGTGCATGGATGTGTACACACCCAAGTCATCAAGACACCATCACACGGATGGCGC<br>CTACTTCGGCACTGGTTTCCCTCACATGCTCTTCATGGTGCATCCCGAGTACCGGC<br>CCAAGAGACCTGCCAACCAGTTTGTGCCCAGGCTCTACGGTTTCAAGATCCATCCG<br>ATGGCCTACCAGCTGCAGCTCCAAGCCGCCAGCAACTTCAAGAGCCCAGTCAAGAC<br>GATTCGC | MRMQLLLLIALSLALVTNSASHHHHHH<br>HHGSSSSEEVSWISWFCGLRGNEFFCE<br>VDEDYIQDKFNLTGLNEQVPHYRQALD<br>MILDLEPDEELEDNPNQSDLIEQAAEM<br>LYGLIHARYILTNRGIAQMLEKYQQGD<br>FGYCPRVYCENQPMLPIGLSDIPGEAM<br>VKLYCPKCMDVYTPKSSRHHHTDGAYF<br>GTGFPHMLFMVHPEYRPKRPANQFVPR<br>LYGFKIHPMAYQLQLQAASNFKSPVKT<br>IR |
| mCD44-Fc-AviTag | atgcgAatgcagctgctgctgctgattgcgctgagcctggcgctggtgaccaacag<br>cactagtCAGATAGACCTTAACGTGACTTGCAGGTACGCTGGCGTTTTTCACGTAG<br>AAAAGAACGGTAGATATAGCATTAGCCGCACAGAAGCTGCGGACCTCTGCCAGGCA<br>TTTAACTCAACGCTGCCAACAATGGACCAAATGAAACTCGCATTGAGTAAAGGTTT<br>TGAAACGTGTCGCTATGGATTTATTGAAGGAAATGTAGTTATTCCCCGGATACACC<br>CAAACGCCATATGTGCCGCGAACCACACCGGAGTTTATATTCTCGTTACATCAAAC<br>ACTAGCCACTACGACACATACTGCTTTAACGCCTCCGCCCCGCCGGAGGAAGATTG<br>TACCTCAGTTACAGACTTGCCCAACTCTTTCGACGGTCCTGTCACAATCACCATCG<br>TAAACCGGGACGGGACAAGGTATTCAAAGAAAGGTGAGTATAGAACACACCAGGAA<br>GATATAGATGCGTCCAATATCATCGACGATGATGTTTCAAGTGGTTCAACAATTGA<br>AAAATCTACGCCTGAGGGTTACATTCTTCATACTTACCTTCCGACAGAGCAGCCTA<br>CAGGCGACCAAGATGACTCCTTTTTTATCAGATCCACCCTTGCAACTATTGCTTCA<br>ACAactagtTCTGGTGGTGGTGGTGAGAATCTGTACTTTCAGAGCTCGGGCGGAGG<br>ATCgggtggaggcgagcccaaatcttgtgacaaaactcacacatgcCCCCCCTGCC | MRMQLLLLIALSLALVTNSTSQIDLNV<br>TCRYAGVFHVEKNGRYSISRTEAADLC<br>QAFNSTLPTMDQMKLALSKGFETCRYG<br>FIEGNVVIPRIHPNAICAANHTGVYIL<br>VTSNTSHYDTYCFNASAPPEEDCTSVT<br>DLPNSFDGPVTITIVNRDGTRYSKKGE<br>YRTHQEDIDASNIIDDDVSSGSTIEKS<br>TPEGYILHTYLPTEQPTGDQDDSFFIR<br>STLATIASTTSSGGGGENLYFQSSGGG<br>SGGGEPKSCDKTHTCPPCPAPELLGGP<br>SVFLFPPKPKDTLMISRTPEVTCVVVD<br>VSHEDPEVKFNWYVDGVEVHNAKTKPR<br>EEQYNSTYRVVSVLTVLHQDWLNGKEY<br>KCKVSNKALPAPIEKTISKAKGQPREP |

| | | |
|---|---|---|
| | CAGCGCCAGAATTGCTGGGCGGACCCAGCGTGTTCCTGTTCCCCCCCAAACCTAAA<br>GACACCCTGATGATCAGCCGAACCCCTGAGGTGACCTGCGTGGTGGTGGACGTGAG<br>CCACGAGGACCCCGAGGTGAAGTTCAACTGGTATGTGGACGGCGTGGAGGTCCACA<br>ATGCCAAAACGAAGCCCAGGGAGGAGCAGTACAACAGCACCTACAGGGTAGTGAGC<br>GTCTTGACCGTGCTGCACCAGGACTGGCTGAACGGCAAGGAATACAAATGCAAGGT<br>CAGCAATAAGGCTCTGCCGGCTCCTATCGAGAAGACAATCAGCAAGGCAAAGGGCC<br>AGCCACGCGAACCGCAGGTGTATACTCTGCCCCCCAGCCGGGACGAGCTGACCAAG<br>AACCAGGTGTCCCTGACCTGTCTGGTGAAAGGCTTCTACCCCAGCGACATCGCTGT<br>GGAGTGGGAGAGTAACGGGCAGCCCGAGAACAACTACAAGACCACGCCTCCTGTGC<br>TGGACAGCGACGGCAGCTTCTTCCTGTATAGCAAGCTCACCGTGGACAAGAGCAGG<br>TGGCAACAGGGCAACGTGTTCAGCTGCTCTGTGATGCACGAGGCCCTGCACAACCA<br>TTACACCCAGAAGAGTCTCAGTCTGAGCCCGGGAAAGGGTGGAGGCGGATCCGGCC<br>TGAACGACATCTTCGAGGCTCAGAAAATCGAATGGCACGAAGGC | QVYTLPPSRDELTKNQVSLTCLVKGFY<br>PSDIAVEWESNGQPENNYKTTPPVLDS<br>DGSFFLYSKLTVDKSRWQQGNVFSCSV<br>MHEALHNHYTQKSLSLSPGKGGGGSGL<br>NDIFEAQKIEWHEG |
| hCD44-<br>Fc-<br>AviTag | atgcgAatgcagctgctgctgctgattgcgctgagcctggcgctggtgaccaacag<br>cactagtCAGATAGACTTGAACATTACTTGCCGCTTCGCCGGTGTGTTTCATGTGG<br>AAAAGAATGGCAGGTACTCTATCTCCAGGACGGAGGCGGCTGATCTCTGCAAGGCA<br>TTCAATTCAACACTTCCGACGATGGCACAGATGGAAAAAGCTCTCAGTATTGGGTT<br>CGAGACGTGCAGATACGGTTTTATTGAGGGCCATGTTGTAATACCCCGCATTCATC<br>CGAATTCTATTTGTGCGGCAAATAATACAGGTGTTTATATCCTTACGTCAAATACG<br>TCTCAGTATGACACCTACTGCTTCAATGCGAGTGCTCCACCGGAAGAAGACTGTAC<br>GTCCGTCACAGATCTGCCTAATGCGTTTGACGGGCCGATCACTATAACGATTGTCA<br>ACAGGGACGGGACTCGATATGTACAAAAGGGGGAATACCGAACAAATCCTGAGGAC<br>ATTTACCCGAGCAATCCTACTGACGACGATGTATCAAGCGGATCCAGTTCCGAACG<br>GTCATCTACCTCCGGGGGGTACATCTTCTATACTTTCTCCACTGTTCACCCCATCC<br>CCGACGAGGATAGTCCATGGATCACTGACTCCACCGACCGAATACCGactagtTCT<br>GGTGGTGGTGGTGAGAATCTGTACTTTCAGAGCTCGGGCGGAGGATCgggtggagg<br>cgagcccaaatcttgtgacaaaactcacacatgcCCCCCCTGCCCAGCGCCAGAAT<br>TGCTGGGCGGACCCAGCGTGTTCCTGTTCCCCCCCAAACCTAAAGACACCCTGATG<br>ATCAGCCGAACCCCTGAGGTGACCTGCGTGGTGGTGGACGTGAGCCACGAGGACCC<br>CGAGGTGAAGTTCAACTGGTATGTGGACGGCGTGGAGGTCCACAATGCCAAAACGA<br>AGCCCAGGGAGGAGCAGTACAACAGCACCTACAGGGTAGTGAGCGTCTTGACCGTG<br>CTGCACCAGGACTGGCTGAACGGCAAGGAATACAAATGCAAGGTCAGCAATAAGGC<br>TCTGCCGGCTCCTATCGAGAAGACAATCAGCAAGGCAAAGGGCCAGCCACGCGAAC<br>CGCAGGTGTATACTCTGCCCCCCAGCCGGGACGAGCTGACCAAGAACCAGGTGTCC<br>CTGACCTGTCTGGTGAAAGGCTTCTACCCCAGCGACATCGCTGTGGAGTGGGAGAG<br>TAACGGGCAGCCCGAGAACAACTACAAGACCACGCCTCCTGTGCTGGACAGCGACG<br>GCAGCTTCTTCCTGTATAGCAAGCTCACCGTGGACAAGAGCAGGTGGCAACAGGGC<br>AACGTGTTCAGCTGCTCTGTGATGCACGAGGCCCTGCACAACCATTACACCCAGAA<br>GAGTCTCAGTCTGAGCCCGGGAAAGGGTGGAGGCGGATCCGGCCTGAACGACATCT<br>TCGAGGCTCAGAAAATCGAATGGCACGAAGGC | MRMQLLLLIALSLALVTNSTSQIDLNI<br>TCRFAGVFHVEKNGRYSISRTEAADLC<br>KAFNSTLPTMAQMEKALSIGFETCRYG<br>FIEGHVVIPRIHPNSICAANNTGVYIL<br>TSNTSQYDTYCFNASAPPEEDCTSVTD<br>LPNAFDGPITITIVNRDGTRYVQKGEY<br>RTNPEDIYPSNPTDDDVSSGSSSERSS<br>TSGGYIFYTFSTVHPIPDEDSPWITDS<br>TDRIPTSSGGGGENLYFQSSGGGSGGG<br>EPKSCDKTHTCPPCPAPELLGGPSVFL<br>FPPKPKDTLMISRTPEVTCVVVDVSHE<br>DPEVKFNWYVDGVEVHNAKTKPREEQY<br>NSTYRVVSVLTVLHQDWLNGKEYKCKV<br>SNKALPAPIEKTISKAKGQPREPQVYT<br>LPPSRDELTKNQVSLTCLVKGFYPSDI<br>AVEWESNGQPENNYKTTPPVLDSDGSF<br>FLYSKLTVDKSRWQQGNVFSCSVMHEA<br>LHNHYTQKSLSLSPGKGGGGSGLNDIF<br>EAQKIEWHEG |
| yeast<br>display<br>vector<br>(no<br>heavy<br>chain) | atgaaggttttgattgtcttgttggctatcttcgctgctttgccattggccttagc<br>tcaaccggttatttctactaccgtcggttccgctgcagaaggctctttggacggtg<br>gatctggcggtggaggttctaagagagaagcttacccatacgatgttccagattac<br>gctGCTAGCGGAGGCGGAGGATCTGGCGGAGGGGGATCAGGCGGAGGCGGCTCCAA<br>CATAGTTATGACCCAGTCCCCTAAATCTATGAGTATGAGCGTAGGTGAGCGAGTAA<br>CTTTGACATGCAAGGCTAGTGAGAACGTGGTAACATATGTCAGTTGGTATCAGCAG<br>AAGCCGGAGCAGTCCCCAAAGTTGCTTATATATGGGGCTAGTAACAGATACACGGG<br>AGTCCCCGACCGCTTCACCGGTAGTGGCTCCGCTACCGACTTTACGTTGACGATTT<br>CCAGCGTTCAAGCGGAAGATTTGGCTGATTATCATTGCGGTCAGGGGAATAGCTAT<br>CCCTACACCTTCGGTGGCGGCACAAAATTGGAGATTAAGgacgcgtcaggtggtgg<br>aggctctggtggcggtggatctggcggtggaggttctgaacaaaagcttatctccg<br>aagaagacttgCAGTTACTTCGCTGTTTTTCAATATTTTCTGTTATTGCTTCAGTT<br>TTAGCACAGGAACTGACAACTATATGCGAGCAAATCCCCTCACCAACTTTAGAATC<br>GACGCCGTACTCTTTGTCAACGACTACTATTTTGGCCAACGGGAAGGCAATGCAAG<br>GAGTTTTTGAATATTACAAATCAGTAACGTTTGTCAGTAATTGCGGTTCTCACCCC<br>TCAACAACTAGCAAAGGCAGCCCCATAAACACACAGTATGTTTTT | MKVLIVLLAIFAALPLALAQPVISTTV<br>GSAAEGSLDGGSGGGGSKREAYPYDVP<br>DYAASGGGGSGGGGSGGGGSNIVMTQS<br>PKSMSMSVGERVTLTCKASENVVTYVS<br>WYQQKPEQSPKLLIYGASNRYTGVPDR<br>FTGSGSATDFTLTISSVQAEDLADYHC<br>GQGNSYPYTFGGGTKLEIKDASGGGGS<br>GGGGSGGGGSEQKLISEEDLQLLRCFS<br>IFSVIASVLAQELTTICEQIPSPTLES<br>TPYSLSTTTILANGKAMQGVFEYYKSV<br>TFVSNCGSHPSTTSKGSPINTQYVF |
| VH1 | gaggttcagctggtggagtctggcggtggcctggtgcagccaggggggctcactccg<br>tttgtcctgtgcagcttctggcttcaacgtctattcttctcctatacactggatgc<br>gtcgggccccgggtaagggcctggaatgggttgcatctatttatccttattctggc<br>tctacttattatgccgatagcgtcaagggccgtttcactataagcgcagacacatc<br>caaaaacacagcctacctacaaatgaacagcctaagagctgaggacagtgccgtct<br>attattgtgctcgctactggccgactggtcattacggttggggtcattggtactgg<br>cattacgatatggactactggggtcaaggaaccctggtcaccgtctccagcGGAGG<br>CGGAGGATCTGGCGGAGGGGGATCAGGCGGAGGCGGCTCCAACATAGTTATGACCC<br>AGTCCCCTAAATCTATGAGTATGAGCGTAGGTGAGCGAGTAACTTTGACATGCAAG<br>GCTAGTGAGAACGTGGTAACATATGTCAGTTGGTATCAGCAGAAGCCGGAGCAGTC<br>CCCAAAGTTGCTTATATATGGGGCTAGTAACAGATACACGGGAGTCCCCGACCGCT<br>TCACCGGTAGTGGCTCCGCTACCGACTTTACGTTGACGATTTCCAGCGTTCAAGCG<br>GAAGATTTGGCTGATTATCATTGCGGTCAGGGGAATAGCTATCCCTACACCTTCGG<br>TGGCGGCACAAAATTGGAGATTAAGgacgcgtcaggtggtggaggctctggtggcg<br>gtggatctggcggtggaggttctTGGTCTCATCCCCAGTTTGAGAAAGGTGGAGGT | EVQLVESGGGLVQPGGSLRLSCAASGF<br>NVYSSPIHWMRRAPGKGLEWVASIYPY<br>SGSTYYADSVKGRFTISADTSKNTAYL<br>QMNSLRAEDSAVYYCARYWPTGHYGWG<br>HWYWHYDMDYWGQGTLVTVSSGGGGSG<br>GGGSGGGGSNIVMTQSPKSMSMSVGER<br>VTLTCKASENVVTYVSWYQQKPEQSPK<br>LLIYGASNRYTGVPDRFTGSGSATDFT<br>LTISSVQAEDLADYHCGQGNSYPYTFG<br>GGTKLEIKDASGGGGSGGGGSGGGGSW<br>SHPQFEKGGGSGGGSGGSSAWSHPQFE<br>K |

| | | |
|---|---|---|
| | TCTGGTGGTGGGTCAGGAGGGAGCAGCGCGTGGTCCCACCCACAATTTGAAAAG | |
| VH2 | ctgatccagttggtgcagtctggacctgagctgaagaagcctggagagacagtcaa<br>gatctcctgcaaggcttctgggtataccttcacaaactatggaatgaactgggtga<br>agcaggctccaggaaagggtttaaagtggatgggctggataaacactgagactggt<br>gagccaacatatgcagatgacttcaagggacggtttgccttctctttggaaacctc<br>tgccagcactgcatatttgcagatcaacaacctcaaaaatgaggacacggctacat<br>atttctgtgcaagacagaggacctacaacggtaatacctttttatgctatggactac<br>tggggtcaaggaaccctggtcaccgtctccagcGGAGGCGGAGGATCTGGCGGAGG<br>GGGATCAGGCGGAGGCGGCTCCAACATAGTTATGACCCAGTCCCCTAAATCTATGA<br>GTATGAGCGTAGGTGAGCGAGTAACTTTGACATGCAAGGCTAGTGAGAACGTGGTA<br>ACATATGTCAGTTGGTATCAGCAGAAGCCGGAGCAGTCCCCAAAGTTGCTTATATA<br>TGGGGCTAGTAACAGATACACGGGAGTCCCCGACCGCTTCACCGGTAGTGGCTCCG<br>CTACCGACTTTACGTTGACGATTTCCAGCGTTCAAGCGGAAGATTTGGCTGATTAT<br>CATTGCGGTCAGGGGAATAGCTATCCCTACACCTTCGGTGGCGGCACAAAATTGGA<br>GATTAAGgacgcgtcaggtggtggaggctctggtggcggtggatctggcggtggag<br>gttctTGGTCTCATCCCCAGTTTGAGAAAGGTGGAGGTTCTGGTGGTGGGTCAGGA<br>GGGAGCAGCGCGTGGTCCCACCCACAATTTGAAAAG | LIQLVQSGPELKKPGETVKISCKASGY<br>TFTNYGMNWVKQAPGKGLKWMGWINTE<br>TGEPTYADDFKGRFAFSLETSASTAYL<br>QINNLKNEDTATYFCARQRTYNGNTFY<br>AMDYWGQGTLVTVSSGGGGSGGGGSGG<br>GGSNIVMTQSPKSMSMSVGERVTLTCK<br>ASENVVTYVSWYQQKPEQSPKLLIYGA<br>SNRYTGVPDRFTGSGSATDFTLTISSV<br>QAEDLADYHCGQGNSYPYTFGGGTKLE<br>IKDASGGGGSGGGGSGGGGSWSHPQFE<br>KGGGSGGGSGGSSAWSHPQFEK |
| VH3 | GTAAAACTCCTTGAATCCGGGGGAGGCCTGGTGCAGCCGGGCGGGAGTCTCAATCT<br>CTCATGTGCAGCTTCAGGCTTCGACTTTTCACGCTACTGGATGAGCTGGGTAAGAC<br>AGGCACCAGGTAAGGGCCAAGAATGGATAGGAGAAATTAACCCCGGAAGCTCTACA<br>ATCAATTATACTCCGTCATTGAAGGACAAGTTCATAATCTCACGCGACAATGCGAA<br>AAATACATTGTATCTTCAAATGAGCAAGGTTAGATCAGAGGATACTGCATTGTATT<br>ACTGTGCTCGCCATGGTAATTATGCGATGGACTATTGGGGCCAGGGAACGCTTGTT<br>ACTGTAAGCAGCGGAGGCGGAGGATCTGGCGGAGGGGGATCAGGCGGAGGCGGCTC<br>CAACATAGTTATGACCCAGTCCCCTAAATCTATGAGTATGAGCGTAGGTGAGCGAG<br>TAACTTTGACATGCAAGGCTAGTGAGAACGTGGTAACATATGTCAGTTGGTATCAG<br>CAGAAGCCGGAGCAGTCCCCAAAGTTGCTTATATATGGGGCTAGTAACAGATACAC<br>GGGAGTCCCCGACCGCTTCACCGGTAGTGGCTCCGCTACCGACTTTACGTTGACGA<br>TTTCCAGCGTTCAAGCGGAAGATTTGGCTGATTATCATTGCGGTCAGGGGAATAGC<br>TATCCCTACACCTTCGGTGGCGGCACAAAATTGGAGATTAAGgacgcgtcaggtgg<br>tggaggctctggtggcggtggatctggcggtggaggttctTGGTCTCATCCCCAGT<br>TTGAGAAAGGTGGAGGTTCTGGTGGTGGGTCAGGAGGGAGCAGCGCGTGGTCCCAC<br>CCACAATTTGAAAAG | VKLLESGGGLVQPGGSLNLSCAASGFD<br>FSRYWMSWVRQAPGKGQEWIGEINPGS<br>STINYTPSLKDKFIISRDNAKNTLYLQ<br>MSKVRSEDTALYYCARHGNYAMDYWGQ<br>GTLVTVSSGGGGSGGGGSGGGGSNIVM<br>TQSPKSMSMSVGERVTLTCKASENVVT<br>YVSWYQQKPEQSPKLLIYGASNRYTGV<br>PDRFTGSGSATDFTLTISSVQAEDLAD<br>YHCGQGNSYPYTFGGGTKLEIKDASGG<br>GGSGGGGSGGGGSWSHPQFEKGGGSGG<br>GSGGSSAWSHPQFEK |
| VH4 | GAATCTGGTCCGGGTTTGGTGAAACCGTCACAATCTCTCTCACTGACCTGCACTGT<br>GACTGGCTACAGCATTACAAGCGATTACGCATGGAACTGGATTCGACAATTTCCGG<br>GCAATAAACTTGAGTGGATGGGTTACATATCCTACTCCGGTTCCACGTCCTACAAC<br>CCTTCTTTGAAATCACGCATATCAATCACCAGAGACACCTCTAAGAACCAATTCTT<br>TCTGCAACTCAATAGTGTGACTACCGAGGACACCGCAACCTACTATTGCGCTAAGA<br>GAATCACAGCCATGGACTATTGGGGACAGGGTACGCTTGTTACTGTTTCCTCAGGA<br>GGCGGAGGATCTGGCGGAGGGGGATCAGGCGGAGGCGGCTCCAACATAGTTATGAC<br>CCAGTCCCCTAAATCTATGAGTATGAGCGTAGGTGAGCGAGTAACTTTGACATGCA<br>AGGCTAGTGAGAACGTGGTAACATATGTCAGTTGGTATCAGCAGAAGCCGGAGCAG<br>TCCCCAAAGTTGCTTATATATGGGGCTAGTAACAGATACACGGGAGTCCCCGACCG<br>CTTCACCGGTAGTGGCTCCGCTACCGACTTTACGTTGACGATTTCCAGCGTTCAAG<br>CGGAAGATTTGGCTGATTATCATTGCGGTCAGGGGAATAGCTATCCCTACACCTTC<br>GGTGGCGGCACAAAATTGGAGATTAAGgacgcgtcaggtggtggaggctctggtgg<br>cggtggatctggcggtggaggttctTGGTCTCATCCCCAGTTTGAGAAAGGTGGAG<br>GTTCTGGTGGTGGGTCAGGAGGGAGCAGCGCGTGGTCCCACCCACAATTTGAAAAG | ESGPGLVKPSQSLSLTCTVTGYSITSD<br>YAWNWIRQFPGNKLEWMGYISYSGSTS<br>YNPSLKSRISITRDTSKNQFFLQLNSV<br>TTEDTATYYCAKRITAMDYWGQGTLVT<br>VSSGGGGSGGGGSGGGGSNIVMTQSPK<br>SMSMSVGERVTLTCKASENVVTYVSWY<br>QQKPEQSPKLLIYGASNRYTGVPDRFT<br>GSGSATDFTLTISSVQAEDLADYHCGQ<br>GNSYPYTFGGGTKLEIKDASGGGGSGG<br>GGSGGGGSWSHPQFEKGGGSGGGSGGS<br>SAWSHPQFEK |
| VH5 | GTGCAGCTGAAACAGTCAGGACCTGGTCTGGTAGCGCCATCACAATCTTTGAGTAT<br>AACGTGCACGGTAAGCGGTTTCAGTCTCACGAGTTACGGAGTGTCCTGGGTCCGAC<br>AGCCCCCAGGCAAGGGACTGGAGTGGTTGGGAGTCATTTGGGGAGATGGCAGTACA<br>AATTACCATTCCGCACTGATTAGTCGATTGTCAATTAGTAAGGATAACAGTAAGTC<br>ACAAGTCTTTCTTAAAATGAACAGCTTGCAAACCGATGACACAGCTATGTATTACT<br>GTGCGAGACATGATTACGCAATGGACTACTGGGGTCAAGGAACACTCGTAACGGTA<br>AGCAGCGGAGGCGGAGGATCTGGCGGAGGGGGATCAGGCGGAGGCGGCTCCAACAT<br>AGTTATGACCCAGTCCCCTAAATCTATGAGTATGAGCGTAGGTGAGCGAGTAACTT<br>TGACATGCAAGGCTAGTGAGAACGTGGTAACATATGTCAGTTGGTATCAGCAGAAG<br>CCGGAGCAGTCCCCAAAGTTGCTTATATATGGGGCTAGTAACAGATACACGGGAGT<br>CCCCGACCGCTTCACCGGTAGTGGCTCCGCTACCGACTTTACGTTGACGATTTCCA<br>GCGTTCAAGCGGAAGATTTGGCTGATTATCATTGCGGTCAGGGGAATAGCTATCCC<br>TACACCTTCGGTGGCGGCACAAAATTGGAGATTAAGgacgcgtcaggtggtggagg<br>ctctggtggcggtggatctggcggtggaggttctTGGTCTCATCCCCAGTTTGAGA<br>AAGGTGGAGGTTCTGGTGGTGGGTCAGGAGGGAGCAGCGCGTGGTCCCACCCACAA<br>TTTGAAAAG | VQLKQSGPGLVAPSQSLSITCTVSGFS<br>LTSYGVSWVRQPPGKGLEWLGVIWGDG<br>STNYHSALISRLSISKDNSKSQVFLKM<br>NSLQTDDTAMYYCARHDYAMDYWGQGT<br>LVTVSSGGGGSGGGGSGGGGSNIVMTQ<br>SPKSMSMSVGERVTLTCKASENVVTYV<br>SWYQQKPEQSPKLLIYGASNRYTGVPD<br>RFTGSGSATDFTLTISSVQAEDLADYH<br>CGQGNSYPYTFGGGTKLEIKDASGGGG<br>SGGGGSGGGGSWSHPQFEKGGGSGGGS<br>GGSSAWSHPQFEK |
